# KIBRA Regulates AMPA Receptor Expression, Synaptic Plasticity, and Memory in an Age-Dependent Manner

**DOI:** 10.1101/2022.02.13.480286

**Authors:** Matthew L. Mendoza, Lilyana Quigley, Thomas Dunham, Lenora J. Volk

## Abstract

The biological mechanisms supporting age-dependent changes in learning and memory remain elusive. While a growing body of human literature implicates KIBRA in memory and neurodevelopmental disorders, KIBRA’s molecular function and contribution to maturation of synaptic function and cognition remain poorly understood. Despite being expressed throughout early postnatal development, germline deletion of KIBRA impairs synaptic plasticity selectively in adult rodents. However, it is unclear whether KIBRA facilitates proper brain maturation necessary for adult plasticity or whether it plays a distinct role in plasticity in the adult brain. Here, using an inducible KIBRA knockout mouse, we demonstrate that acutely deleting KIBRA in adult forebrain neurons impairs both spatial memory and long-term potentiation (LTP). The deficits in LTP correlate with an adult-selective decrease in extrasynaptic AMPA receptors under basal conditions. We also identify a novel role for KIBRA in LTP-induced AMPAR upregulation. In contrast, acute deletion of KIBRA in juvenile forebrain neurons did not affect LTP and had minimal effects on basal AMPAR expression. These data suggest that KIBRA serves a unique role in adult hippocampal function through regulation of basal and activity-dependent AMPAR proteostasis that supports synaptic plasticity.

**Significance Statement:** Synaptic plasticity supported by trafficking of postsynaptic AMPA receptors is a conserved mechanism underlying learning and memory. The nature and efficacy of learning and memory undergo substantial changes during childhood and adolescent development, but the mechanisms underlying this cognitive maturation remain poorly understood. Here, we demonstrate that the human memory- and neurodevelopmental disorder-associated gene KIBRA facilitates memory and hippocampal synaptic plasticity selectively in the adult hippocampus. Furthermore, we show that selective loss of KIBRA from adult but not juvenile neurons reduces expression of extrasynaptic AMPA receptors and prevents LTP-induced increases in AMPAR expression. Overall, our results suggest that KIBRA participates in cellular and molecular processes that become uniquely necessary for memory and synaptic plasticity in early adulthood.

## Introduction

Learning and memory are dynamic processes that evolve throughout the lifespan of an organism. The early postnatal period (1-2 years in humans, 1-3 weeks in rodents) is accompanied by rapid growth in synaptic density and maturation of molecular composition at synapses (Semple et al., 2013; Lohmann and Kessels, 2014). The closing of this period coincides with the late postnatal emergence of hippocampal-dependent memory (Dumas, 2005; Alberini and Travaglia, 2017) and accurate representation of spatial and sequential information in the hippocampal network (Farooq and Dragoi, 2019). Substantial effort has focused on understanding the synaptic changes that accompany early postnatal development. However, hippocampal-dependent memory in rodents continues to mature toward adult-like performance throughout adolescence (Alberini and Travaglia, 2017), and much less is known about changes in synaptic plasticity mechanisms across this critical developmental time period. Childhood and adolescent development also coincide temporally with the onset of genetically overlapping neurodevelopmental disorders (NDDs) that have in common some form of synaptic dysfunction.

Numerous studies demonstrate that common variants of *KIBRA* (expressed in <u>Ki</u>dney and <u>BRA</u>in; a.k.a. *WWC1*) associate with human memory performance (Papassotiropoulos et al., 2006; Almeida et al., 2008; Schaper et al., 2008; Bates et al., 2009; Preuschhof et al., 2009; Vassos et al., 2010; Yasuda et al., 2010; Kauppi et al., 2011; Pawlowski and Huentelman, 2011; Milnik et al., 2012; Duning et al., 2013; Muse et al., 2014; Vyas et al., 2014; Rovira et al., 2016). Supporting a role in mnemonic function, reduction or overexpression of KIBRA impairs memory in rodents (Makuch et al., 2011; Vogt-Eisele et al., 2014; Heitz et al., 2016), and KIBRA expression is enriched in the human and rodent hippocampus (Allen-Institute_Brain-Atlas; Papassotiropoulos et al., 2006), a brain structure critical for memory formation. Consistent with its effects on memory, deletion or overexpression of KIBRA impairs synaptic plasticity (Long-Term Potentiation, LTP and Long-Term Depression, LTD) in rodents (Makuch et al., 2011; Heitz et al., 2016) and blocks associative long term facilitation in *Aplysia* (Hu et al., 2017). However, despite robust early postnatal expression (Johannsen et al., 2008) (Fig 1A), constitutive deletion of KIBRA impairs synaptic plasticity selectively in young adult (2-3.5 months-old) but not juvenile (3-4 week-old) mice (Makuch et al., 2011). It is currently unclear if this distinction results from juvenile resilience and/or adult vulnerability to loss of KIBRA, or whether KIBRA is required for maturation of synaptic function during adolescence. Interestingly, *KIBRA* polymorphisms associate with developmentally-regulated disorders of complex cognition, including schizophrenia (SCZ) (Kos et al., 2016) and Tourette (Willsey et al., 2017), which also show adolescent or young adult onset. KIBRA is a postsynaptically-localized scaffolding protein, and a large proportion of neuronally-expressed KIBRA binding partners also associate with neurodevelopmental disorders including SCZ (Hakak et al., 2001; Dev and Henley, 2006; Lauriat et al., 2006; Fromer et al., 2016), bipolar disorder (BPD) (Hou et al., 2016), and/or autism spectrum disorder (ASD) (Voineagu et al., 2011; Parikshak et al., 2016).

**Figure 1.**
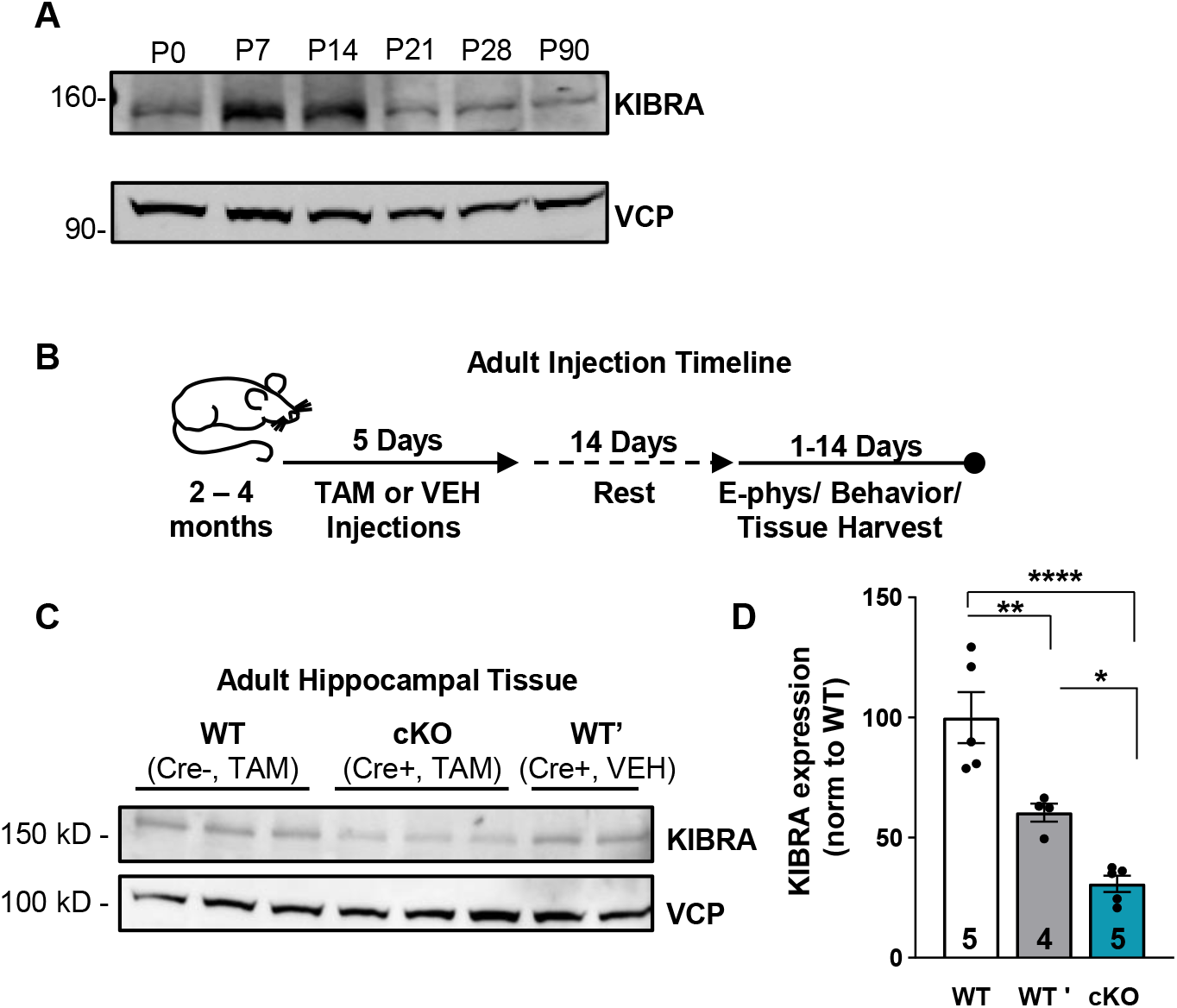
**Tamoxifen treatment acutely reduces hippocampal KIBRA expression in the adult brain**. (**A**) Developmental expression profile of KIBRA in the hippocampus of WT mice (**B**) Tamoxifen injection schedule for adult Kibra^floxed/floxed^:CaMK2α Cre^ERT2^ mice. WT (Cre-negative, tamoxifen injected), WT’ (Cre-positive, vehicle injected), and cKO (Cre-positive, tamoxifen injected) mice received 2 injections (100mg/kg I.P.) per day for 5 days. Each mouse was given 14 days to recover from injections before experiments and tissue collection. Injections and experiments were performed after each mouse turned 2 months of age and before 4.5 months of age. (**C**) Hippocampal CA1 tissue was isolated from Kibra^floxed/floxed^:CaMK2α Cre^ERT2^ mice following tamoxifen or vehicle injections. Endogenous KIBRA protein levels were assessed using an anti- KIBRA antibody. (**D**) Quantification normalized to VCP: One-way ANOVA with Holm Sidak’s multiple comparisons test. (WT, 100 ± 10.58%; WT’, 60.44± 3.71%; cKO, 30.88 ± 3.39%). *p<0.05, **p<0.01, ****p<0.0001. Data plotted as mean ± SEM, n = number of animals, indicated on each bar.

**Table 1.1:**
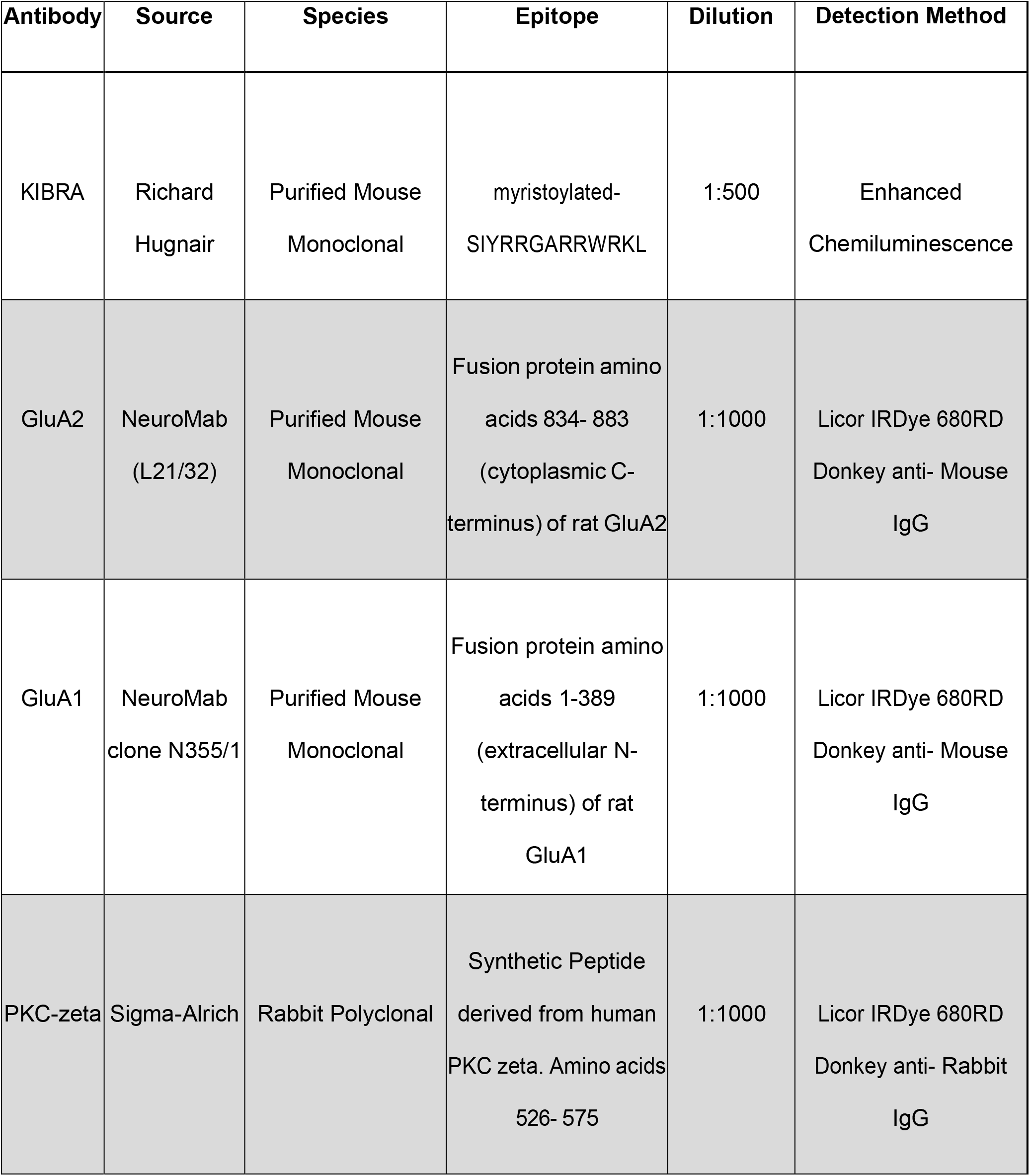
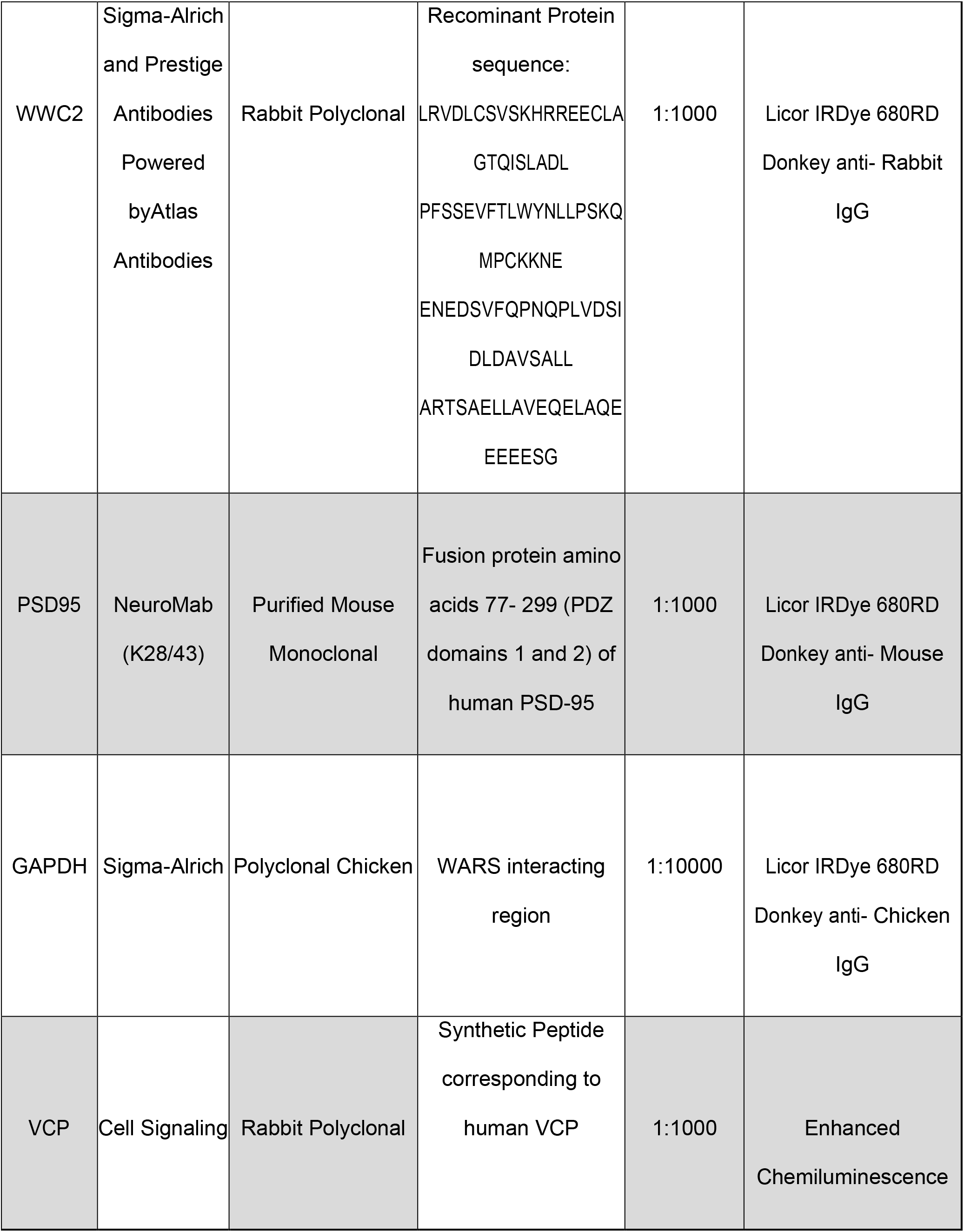
Primary and Secondary Antibodies

Activity-dependent changes in postsynaptic AMPAR (α-amino-3-hydroxy-5-methyl-4- isoxazolepropionic acid receptor) concentration, localization, and function are highly conserved expression mechanisms of synaptic plasticity (Shepherd and Huganir, 2007; Huganir and Nicoll, 2013; Diering and Huganir, 2018). In neurons, KIBRA concentrates at excitatory synapses (Makuch et al., 2011; Heitz et al., 2016) where it regulates trafficking of AMPA-type glutamate receptors (Makuch et al., 2011; Heitz et al., 2016; Tracy et al., 2016). While the specific mechanisms by which KIBRA regulates AMPAR trafficking are unknown, in non-neuronal cells KIBRA mediates recycling of endocytosed cargo through Rab11 positive recycling endosomes (Traer et al., 2007). KIBRA is a scaffolding protein with multiple protein and lipid interaction domains (Zhang et al., 2014). The KIBRA interactome includes regulators of endosomal trafficking (Traer et al., 2007; Makuch et al., 2011; Fukuda et al., 2019), actin (Kremerskothen et al., 2003; Kremerskothen et al., 2006; Duning et al., 2008; Rocca et al., 2008; Makuch et al., 2011) and exocytic pathways (Rosse et al., 2009; Makuch et al., 2011; Song et al., 2019), suggesting that KIBRA may regulate endosomal sorting of AMPARs.

Additionally, loss of KIBRA reduces the basal expression of multiple binding partners including PKMζ (Vogt-Eisele et al., 2014; Hu et al., 2017; Ferguson et al., 2019), Lats1/2 (Xiao et al., 2011), and Rab27a (Song et al., 2019). However, it is currently unclear if KIBRA regulates AMPA proteostasis, or if KIBRA’s regulation of AMPA receptors changes with age.

Here, using inducible knockout to acutely deplete KIBRA from CaMKIIα+ forebrain neurons in adult or juvenile mice, we show that acute reduction of KIBRA causes depletion of extrasynaptic pools of AMPARs and impairs LTP selectively in the adult hippocampus. We identify a novel role for KIBRA in maintaining LTP-induced increases in AMPAR expression in adult mice, and show that KIBRA expression in neurons is required to maintain long-term memory.

## Results

### Acute reduction of KIBRA in neurons reduces long-term potentiation in the adult hippocampus

Synaptic plasticity is impaired in adult (2-3.5 months old), but not juvenile (3-4 week old) constitutive KIBRA KO mice (Makuch et al., 2011). This is not due to lack of KIBRA expression at juvenile ages in WT mice. In fact, KIBRA expression is higher in early postnatal mice than in adults at the mRNA (Johannsen et al., 2008) and protein (Fig. 1A) level. Whether this adult-selective impairment in synaptic plasticity results from a requirement for KIBRA during adolescent development, juvenile resilience/adult vulnerability to loss of KIBRA (juvenile-selective ability to compensate), and/or a unique requirement for KIBRA in adult forms of synaptic plasticity is unknown. To determine if synaptic plasticity in adult neurons is vulnerable to loss of KIBRA, and to disambiguate this from a role in synapse or circuit development, we crossed conditional *Kibra^Floxed/Floxed^(Makuch et al., 2011)* mice with CaMKII CreER^T2^ (Erdmann et al., 2007) mice that express tamoxifen-inducible Cre recombinase (a fusion protein containing Cre recombinase and the ligand-binding domain of the estrogen receptor) under the CaMKIIα promoter. This generated mice in which genetic deletion of KIBRA is restricted to primarily excitatory forebrain neurons and is temporally controlled via tamoxifen injections, thus providing age, cell-type, and region-selective control of gene expression. Adult (2-4 months old) *Kibra^Fl/Fl^*:CaMKIICreER^T2+^ mice (KIBRA cKO) that received tamoxifen injections showed approximately 70% reduction in KIBRA protein expression (Fig. 1 B-D). We interpret the residual KIBRA expression to be largely due to the fact that while Cre is primarily expressed in excitatory neurons, KIBRA is also expressed in a subset of inhibitory neurons and astrocytes (Allen-Institute_Cell-Types; Cembrowski et al., 2016; Habib et al., 2016). We also observe a ∼40% reduction in KIBRA expression in vehicle-treated Cre-positive adult mice (WT’, *Kibra^Fl/Fl^*:CaMKIICreER^T2+^) compared to tamoxifen-treated Cre-negative mice (WT, *Kibra^Fl/Fl^*:CaMKIICreER^T2-^) mice, indicating low-levels of spontaneous Cre activation (Fig. 1C-D). Importantly, because Cre expression is under the control of the CamKIIα promoter (Erdmann et al., 2007), we observe a selective decrease in KIBRA in adult WT’ mice, with no changes in KIBRA during early postnatal development (Fig. 6, no reduction in KIBRA in juvenile WT’ mice)..

**Figure 6.**
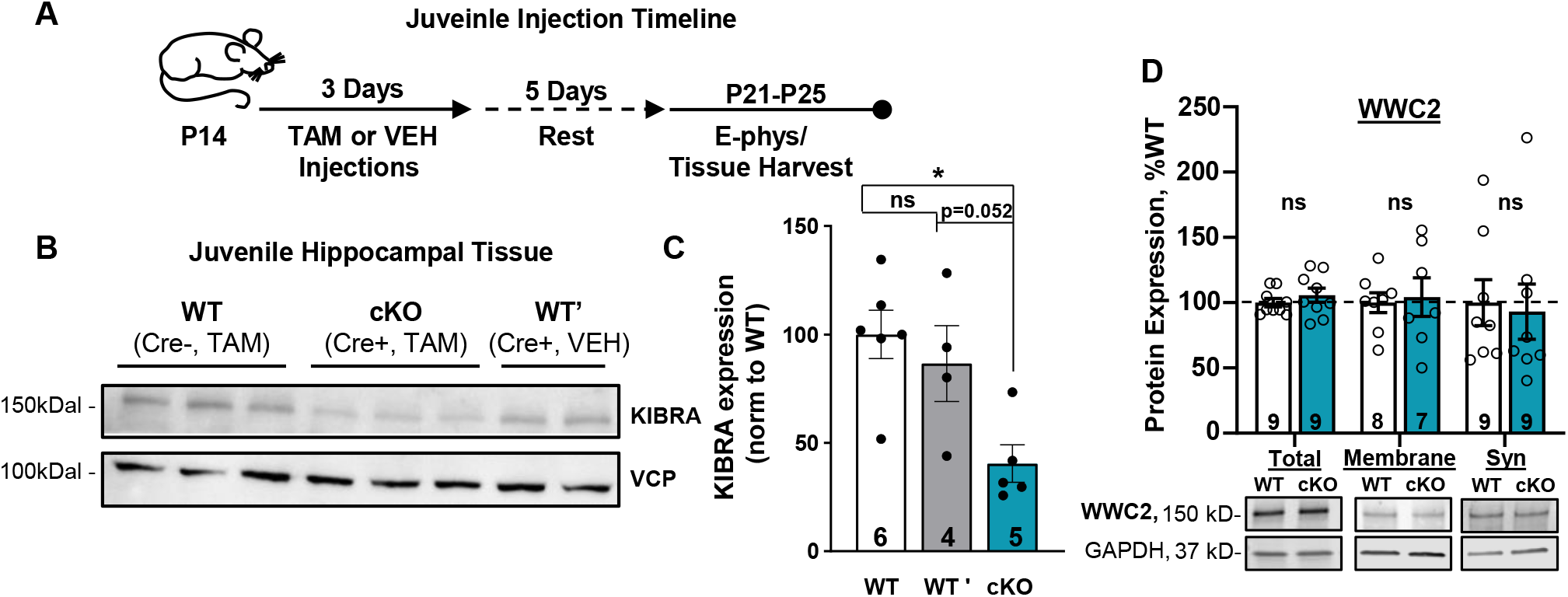
**Tamoxifen treatment acutely reduces KIBRA expression in the juvenile hippocampus**. (**A**) Tamoxifen injection schedule for juvenile Kibra^floxed/floxed^:CaMK2α Cre^ERT2^ mice. WT (Cre-negative, tamoxifen injected), WT’ (Cre-positive, vehicle injected), and cKO (Cre-positive, tamoxifen injected) mice received 1 injection (100mg/kg I.P.) per day for 3 days. Each mouse was given 5 days to recover from injections before experiments and tissue collection. Experiments were performed between P21-P25. (**B**) Hippocampal CA1 tissue was isolated following tamoxifen or vehicle injections. Endogenous KIBRA protein levels were assessed using an anti- KIBRA antibody. (**C**) Quantification normalized to VCP (one-way ANOVA, p = 0.010, post hoc comparisons shown in figure). WT, 100 ± 11%; WT’, 87 ± 17%; cKO, 40 ± 9%. (**D**) Acute reduction of KIBRA in the juvenile brain does not affect expression of the KIBRA homolog WWC2 (unpaired t-test (total and membrane) or Mann-Whitney test (synaptic) corrected for multiple comparisons). Total, WT=100 ± 3%, cKO=106 ± 5%; membrane, WT=100 ± 7%, cKO=104 ± 14%; synaptic, WT=100 ± 17%, cKO=93 ± 21%. *p<0.05, 0.05 <#p < 0.1, n.s. p>0.1. Data plotted as mean ± SEM, n = number of animals, indicated on each bar.

We first determined if reduction of KIBRA alters basal synaptic transmission or presynaptic release probability in adult cKO mice. Using extracellular field potential (fEPSP) recordings of Schaffer Collateral synapses in acute hippocampal slices, we observe that the level of KIBRA expression in adult mice does not influence input-output relationship (Fig. 2A-C), indicating that basal synaptic transmission is unaffected by KIBRA depletion. Next, we examined whether reducing KIBRA influenced the presynaptic release probability of Schaffer Collateral CA1 synapses. We observe no difference in paired-pulse facilitation at CA1 synapses, indicating that acutely reducing KIBRA in the adult brain has no impact on excitatory presynaptic release probability (Fig. 2D). Collectively, these data argue that acute reduction of KIBRA in the adult brain does not impact basal synaptic transmission, consistent with data from adult whole-body, constitutive KIBRA knockout (KO) mice (Makuch et al., 2011).

**Figure 2.**
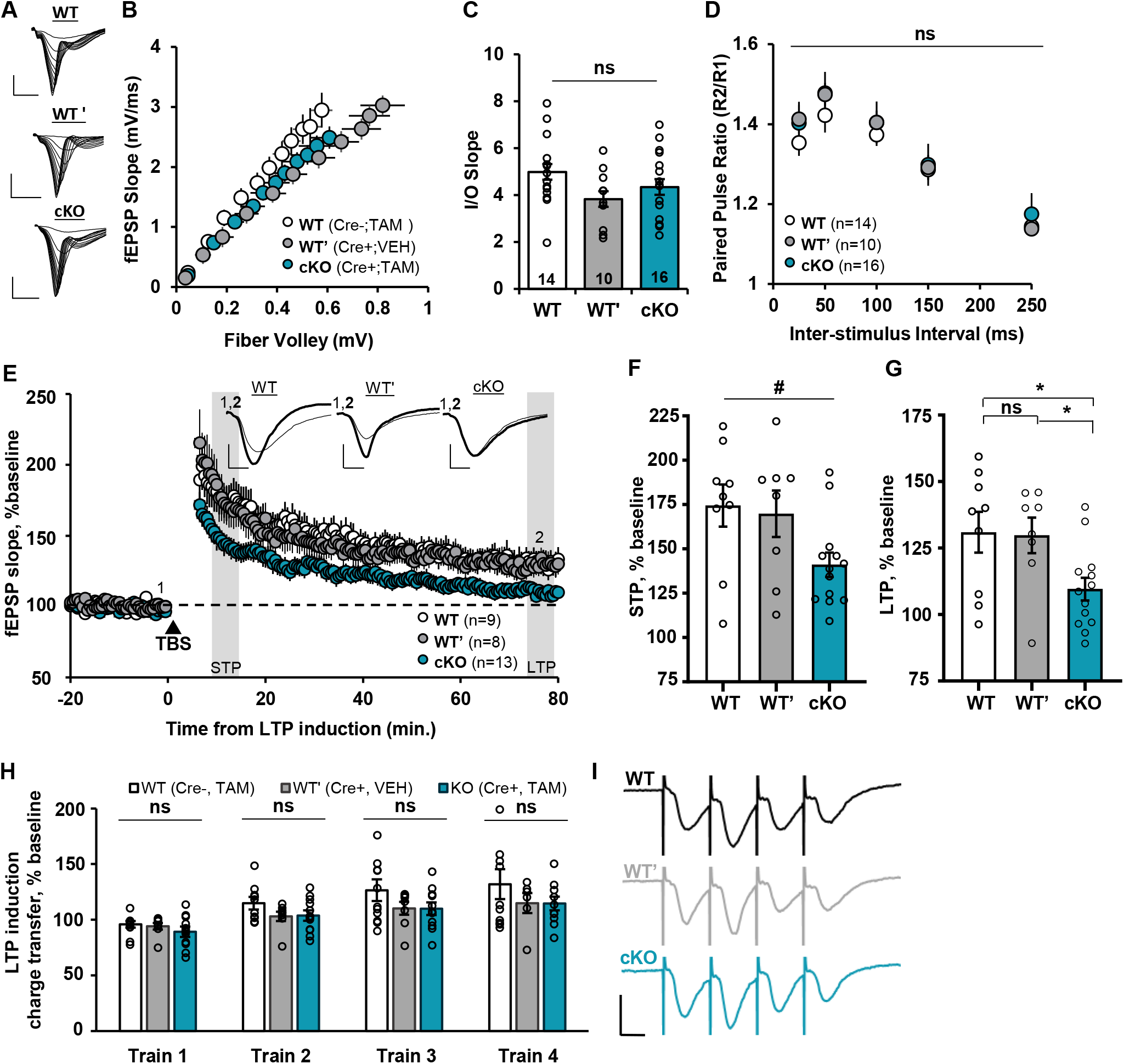
KIBRA acutely regulates hippocampal long-term potentiation without influencing basal synaptic transmission in the adult brain. (A) Representative traces from input-output curves, scale bars = 2mV/5ms, (B) Summary data from adult input-output analysis, (C) Slopes of individual input-output curves: WT, average I-O slope 4.98 ± 0.41 ms^-1^; WT’, average I-O slope 3.83 ± 0.36 ms^-1^; cKO, I-O slope 4.34 ± 0.39 ms^-1^. No difference observed across experimental conditions (one-way ANOVA, n.s.). (D) Paired pulse facilitation (fEPSP slope response 2/fEPSP slope response1) is not altered in adult KIBRA cKO mice (repeated measures (RM) two-way ANOVA, n.s. genotype X inter-stimulus interval interaction, main effect of genotype, and multiple comparisons at all inter-stimulus intervals). (E) Hippocampal LTP induced by four trains of theta-burst stimulation is impaired after acute reduction of KIBRA in the adult brain. Scale bars= 0.25mV/5ms. (F) Trend towards decreased STP in adult KIBRA cKO mice (one-way ANOVA, p = 0.063). 5min average at grey bar in E, WT, 174 ± 12 %; WT’, 170 ± 13 %; cKO, 141 ± 7. (G) LTP magnitude measured at 75-80 mins post LTP induction is decreased after acute reduction of KIBRA in the adult hippocampus (one-way ANOVA, p = 0.021). 5min average at grey bar in E, WT, 130 ± 8%; WT’, 130 ± 7 %; cKO, 110 ± 4%. (H) LTP induction is not impaired in adult KIBRA cKO mice (one-way ANOVA, n.s. effect of genotype, genotype x train interaction, and multiple comparisons at all inter-stimulus intervals). LTP induction measured as charge transfer (Area Under Curve, AUC) during each TBS train (40 fEPSPs per train, normalized to baseline fEPSP). (I) Sample traces from LTP induction quantified in H. Shown is the first burst of the first train from one example recording (not averaged), Scale bar = 1.5mV/50 ms. All summary data presented as mean ± SEM. *p<0.05, # 0.5<p<0.1, n.s. (not significant) = p>0.1. B-D, WT, n=14 slices from 6 mice; WT’, n = 10 slices from 3 mice; cKO, n = 16 slices from 6 mice. E-H, WT, n=9 slices from 5 mice; WT’, n = 8 slices from 3 mice; cKO, n = 13 slices from 5 mice.

To determine if KIBRA is specifically required for synaptic plasticity in the adult brain independent of potential neurodevelopmental confounds, we utilized our inducible KIBRA cKO mice to examine LTP in adult mice. Acutely deleting KIBRA in the adult brain significantly reduces LTP expression (Fig. 2E,G). We also observe a trend towards decreased short-term potentiation (Fig. 2E,F), a process that requires lateral diffusion of surface-localized extrasynaptic AMPARs (Penn et al., 2017). Importantly the WT’ group shows LTP and STP similar to WT mice (Cre-negative tamoxifen-treated group, Fig. 2E-G). In contrast to LTP expression, KIBRA deletion had no effect on charge transfer during the LTP induction stimulus (Theta-burst stimulation; TBS; see methods) (Fig. 2H,I). These data highlight the importance of KIBRA for expression of adult hippocampal synaptic plasticity.

### Acute reduction of KIBRA in adult neurons reduces basal expression of extrasynaptic AMPARs in the hippocampus

AMPAR trafficking into and out of the synapse is a core mechanism driving the postsynaptic expression of synaptic plasticity (Choquet, 2018; Diering and Huganir, 2018). Neurons maintain distinct pools of AMPARs at excitatory synapses, including a synaptically localized pool which mediates neuronal communication, as well as extra- synaptic surface and intracellular pools that can be rapidly mobilized and inserted into the synapse to facilitate synaptic potentiation (Penn et al., 2017). KIBRA forms a complex with AMPARs (Makuch et al., 2011; Heitz et al., 2016) and has been shown to prevent degradation of multiple binding partners (Xiao et al., 2011; Vogt-Eisele et al., 2014; Hu et al., 2017; Ferguson et al., 2019; Song et al., 2019), thus we hypothesized that loss of KIBRA disrupts AMPAR proteostasis and depletes AMPARs from extrasynaptic pools required for LTP. To test this hypothesis, we performed subcellular fractionation on hippocampal area CA1 tissue from KIBRA cKO and WT mice and examined total, membrane-localized, and synaptically-localized (postsynaptic density-associated) AMPARs (Chiu et al., 2017). Acute reduction of KIBRA in adult neurons decreases total expression of the AMPAR subunits GluA2 and GluA1 (Fig. 3A,B) supporting a role for KIBRA in AMPAR proteostasis. Consistent with prior work, we also observe a reduction in PKMζ (WT, 100 ± 2%; cKO 74 ± 2%, p < 0.001), a brain-specific kinase that interacts with KIBRA (Buther et al., 2004; Vogt-Eisele et al., 2014; Ferguson et al., 2019). Acute reduction of KIBRA also decreased the basal expression of KIBRA homolog and binding partner WWC2 (Fig. 3G,H). Total PSD95 expression is unchanged in cKO mice, indicating acute KIBRA deletion does not broadly and non-selectively reduce synaptic protein expression (Fig 3A,B). GluA2 levels were also reduced in the membrane fraction of KIBRA cKO mice (Fig. 3C-D), supporting the hypothesis that KIBRA facilitates maintenance of extrasynaptic AMPAR pools. GluA1 expression was not decreased in the KIBRA cKO membrane fraction, suggesting that under basal conditions GluA1 is depleted from intracellular stores. Synaptic expression of AMPARs was unchanged (Fig. 3E-F) in KIBRA cKO mice, consistent with our observation of normal basal synaptic transmission in these mice (Fig. 2A-D).

**Figure 3.**
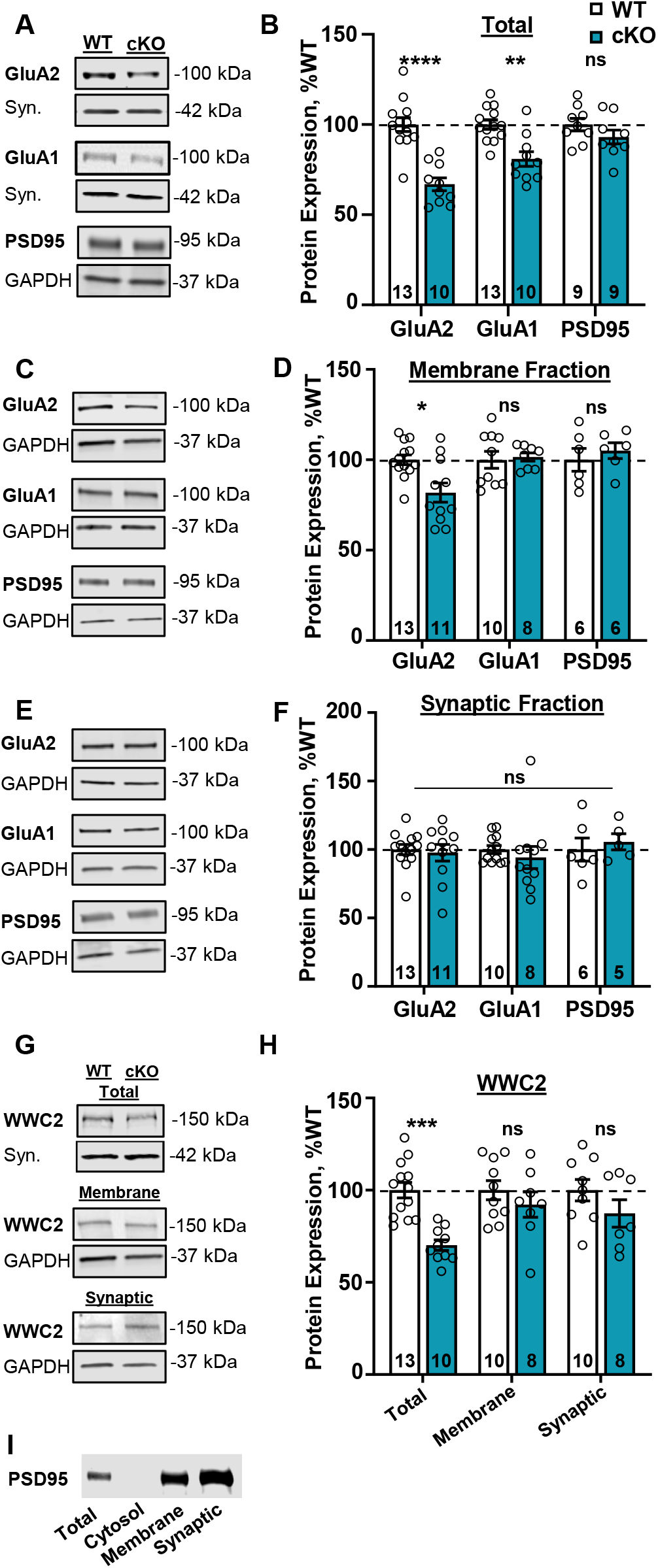
KIBRA acutely regulates the basal expression of extrasynaptic AMPA receptors in the adult hippocampus. (**A,C,E,G)** Representative western blot images from sub-region CA1 of the adult hippocampus. Samples are normalized to loading control and quantified as % WT in **B,D,F,H** (see methods for details). (**B**) Acute reduction of KIBRA in the adult brain decreases the total expression GluA2 and GluA1, but not the excitatory synaptic scaffold PSD-95 (unpaired t-tests, corrected for multiple comparisons). GluA2, WT= 100 ± 3%, cKO=67 ±3; GluA1, WT=100 ± 2%, cKO=80 ± 4%; PSD95, WT=100 ± 3%, cKO=93 ± 3%. (**D**) Acute reduction of KIBRA in the adult brain decreases expression of membrane-associated GluA2 (unpaired t-tests with Welch’s correction, corrected for multiple comparisons). GluA2, WT=100 ± 2%, cKO=83 ± 5%; GluA1, WT=100 ± 4%, cKO=102 ± 2%; PSD95, WT=100 ± 6%, cKO=105 ± 4%. (**F**) Acute reduction of KIBRA in the adult brain does not alter basal expression of synaptic AMPA receptors (unpaired Mann-Whitney tests, corrected for multiple comparisons). GluA2, WT=100 ± 3%, cKO=98 ± 6%; GluA1, WT=100 ± 2%, cKO=94 ± 8%; PSD95, WT=100 ± 8%, cKO=106 ± 5%. (**H**) Decrease in total expression of the KIBRA homolog WWC2 following acute reduction of KIBRA in the adult brain (unpaired t-tests, corrected for multiple comparisons). Total, WT=100 ± 4%, cKO=70 ± 2%; membrane, WT=100 ± 5%, cKO=92 ± 6%; synaptic, WT=100 ± 5%, cKO=87 ± 7%. (**I**) Representative western blot with equal protein loaded for total, cytosolic, membrane and synaptic fractions, demonstrating depletion of the postsynaptic scaffold PSD95 from the cytosolic fraction and enrichment in the membrane fraction with further enrichment in the synaptic fraction. Data shown as mean ± SEM, n on bar graphs = number of animals.

### KIBRA is necessary for LTP-induced upregulation of AMPARs in the adult hippocampus

One mechanism by which neural activity can facilitate synaptic potentiation is through rapid *de novo* synthesis of AMPARs (Nayak et al., 1998; Matsuo et al., 2008; Tushev et al., 2018). While previous studies have examined the role of KIBRA in activity-dependent AMPAR trafficking via cell-wide pharmacological stimulation in cultured neurons with exogenously-expressed tagged AMPRs (Makuch et al., 2011; Heitz et al., 2016), it is unknown whether KIBRA regulates expression of endogenous AMPARs in response to physiological plasticity-inducing stimuli. Therefore, we examined whether KIBRA influences LTP-induced increases in endogenous AMPAR expression in the adult hippocampus. Acute hippocampal slices were subjected to basal stimulation (0.033Hz) or TBS LTP, which we have previously shown to be a protein synthesis-dependent form of synaptic plasticity (Volk et al., 2013), followed by microdissection of the stimulated area in CA1 at 30 or 120 minutes after LTP induction (Fig. 4A). LTP induced an increase in GluA2 (Fig. 4B,C) and GluA1 (Fig. 4D,E) expression in WT but not KIBRA cKO mice. These data suggest that KIBRA not only prevents basal degradation of AMPARs, but is also necessary to maintain activity-induced increases in AMPAR expression after LTP.

**Figure 4.**
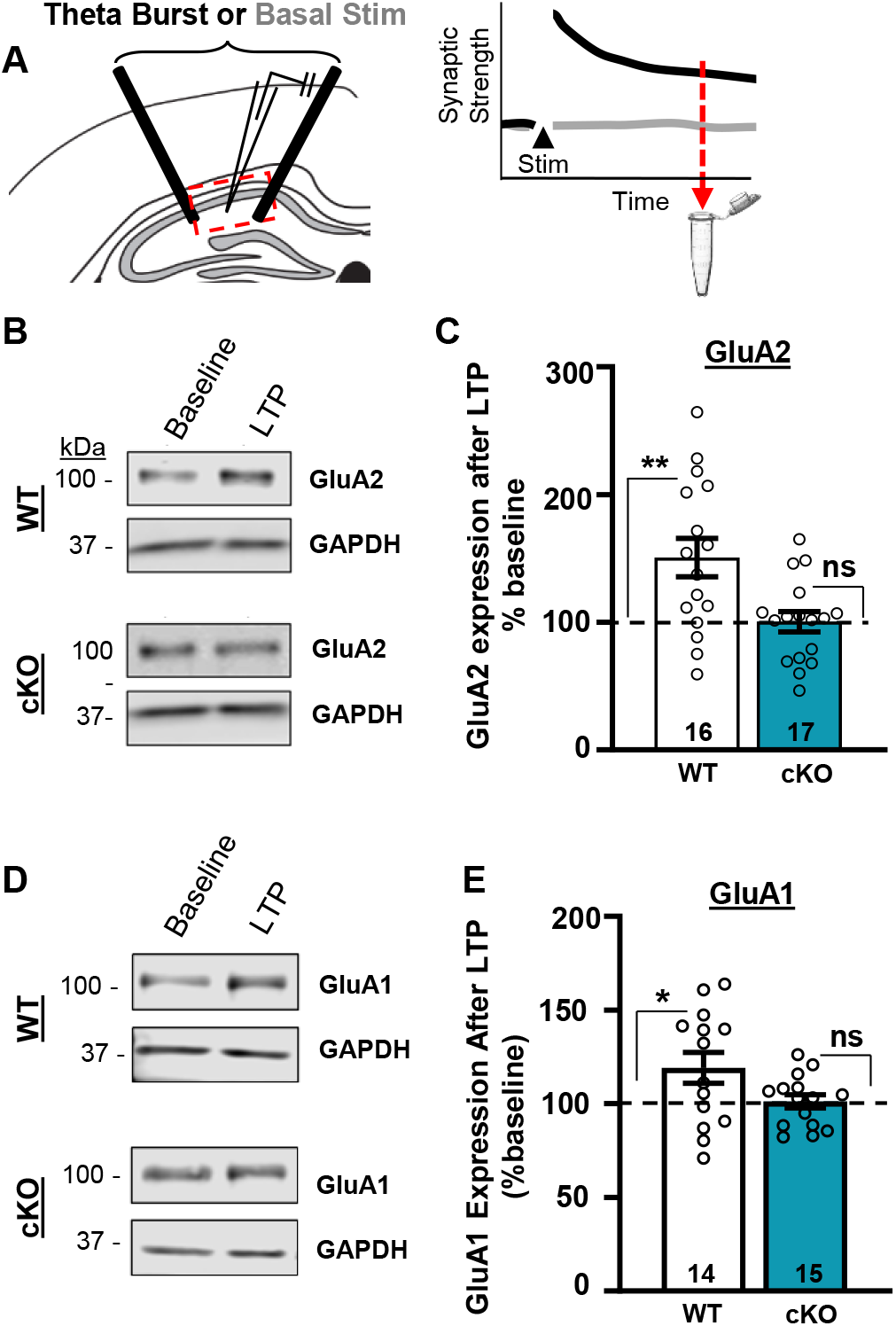
KIBRA is required for LTP-induced increase in AMPAR expression. (**A**) Experimental design. Transverse hippocampal slices were collected from adult mice following basal stimulation (0.033 Hz) or LTP (TBS). The stimulated region of CA1 was microdissected 30 or 120 mins after LTP or basal stimulation. Data from both time points was combined as no difference in AMPAR induction was observed between 30 and 120min post LTP. (**B,D**) Representative GluA2 and GluA1 immunoblot images after baseline stimulation or LTP from WT (cre-negative tamoxifen treated) or KIBRA cKO adult mice (cre-positive tamoxifen treated). LTP increases GluA2 (**C**) and GluA1 (**E**) expression in WT but not KIBRA cKO mice. Data plotted as % increase over baseline stimulation, mean ± SEM (GluA2, WT mean= 151 ± 15 %, cKO = 100 ± 8%; GluA1, WT mean= 119 ± 8%, cKO = 101 ± 4%). Number of slices is indicated on each bar. One sample t-test, *p< 0.05, **p< 0.01 .

### Acute reduction of KIBRA in forebrain neurons impairs spatial memory in adult mice

KIBRA expression is highest in excitatory neurons, but it is also present in a subset of inhibitory neurons and glial cells (Allen-Institute_Cell-Types; Cembrowski et al., 2016; Habib et al., 2016). The cell-type(s) in which KIBRA is required to support memory function are not known. Given its critical role in hippocampal LTP, we next determined the mnemonic role of KIBRA in CamKIIα+ forebrain neurons by assessing spatial memory in adult KIBRA CamKIIα Cre-KO ^ERT2^ mice. Mice were trained to navigate to a rewarded escape box (target) on a modified Barnes Maze (Fig 5A). Training consisted of four trials per day over four consecutive days. We quantified the latency to reach the target on the first trial of each day as a measure of memory retention across training sessions. WT mice showed significant improvement from Day 1 to Day 2, with continued improvement thereafter (Fig 5B). However, KIBRA cKO littermates did not show significant improvement until Day 4, suggesting delayed memory retention during training (Fig 5B). When all trials across a day were combined, KIBRA cKO mice were not statistically different from WT littermates (Fig 5C). Thus, learning in KIBRA cKO mice was slowed but not abolished, and the impaired performance on trial 1 of Days 2 and 3 (Fig 5C) was not due to reduced levels of learning the prior day. We next tested long-term memory via a single probe trial (no target box) 7 days after the final training day (Fig 5D). Whereas WT mice showed significant preference for the target quadrant, KIBRA cKO mice had similar occupancy for the target and neighboring quadrants (Fig 5E), indicating an impairment in long-term spatial memory. KIBRA cKO mice also showed fewer target entries within the first five entries (Fig 5F). Thus, KIBRA expression in CaMKIIα+ forebrain neurons appears necessary for maintenance of long-term spatial memory. Overall movement (Fig 5G, H), time spent in the center of the maze (Fig.5I), and weight reduction during time-restricted feeding (Fig 5J) were not different between genotypes, indicating that memory deficits in KIBRA cKO mice were not due to changes in locomotion, anxiety-like behaviors on the apparatus, or different responses to food restriction.

**Figure 5.**
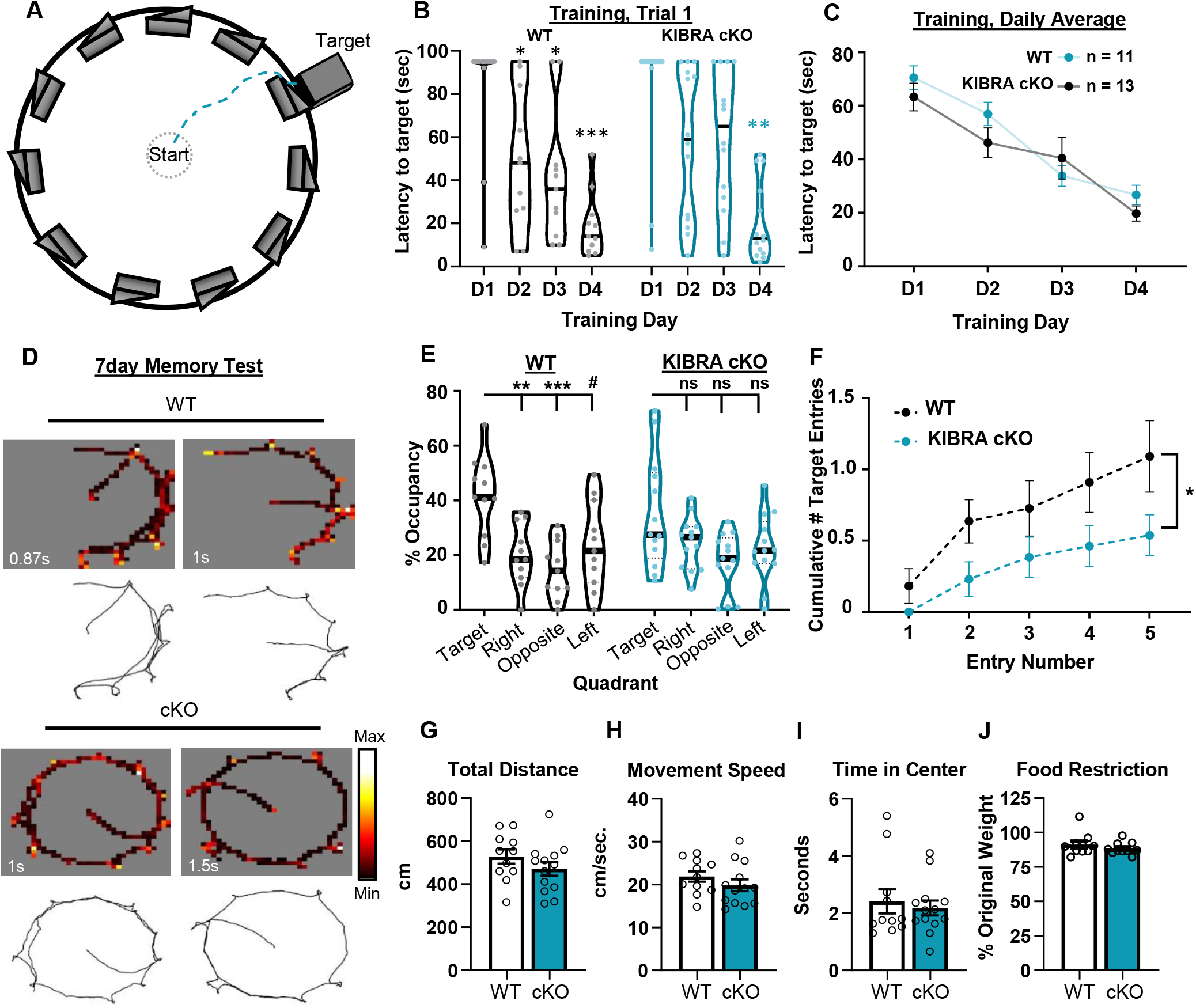
Acute loss of KIBRA in neurons impairs memory in adult mice. (**A**) Schematic of modified Barnes Maze arena. The target location is a covered box hidden from view by one of the identical walls positioned around the exterior of the maze. Training consisted of 4 trials per day over four days. (**B**) KIBRA mice show delayed memory acquisition during training, assessed by comparing latency to reach the target box on the first trial of day 2,3,and 4 to the first trial on day 1 (Welch’s RM one-way ANOVA, cKO p = 0.0013, WT p = 0.0006, post-hoc comparisons shown in figure). Lines in violin plots = median. (**C**) KIBRA cKO mice exhibit grossly normal learning as assessed by latency to reach the target box averaged across all 4 trials for each training day (RM two-way ANOVA, n.s. effect of genotype, genotype x day interaction, or WT vs KO post hoc comparison for any day). n = number of mice: WT, 6M + 5F; KIBRA cKO 8M +6F. (**D**) Example maze occupancy (top) and trajectory (bottom) plots during memory retention probe test from two example mice of each genotype. ‘Max’ time for occupancy scale is indicated at the bottom left corner of each plot. Target location is as depicted in panel A. (**E**) Percent occupancy per zone during memory retention (probe) test 7 days after the final training session. KIBRA cKO mice fail to show preference for target quadrant (Welch’s RM one-way ANOVA, cKO p = 0.0913, WT p = 0.0137, post- hoc comparisons shown in figure) . Lines in violin plots = median. . (**F**) Decreased memory exhibited by KIBRA cKO mice shown by the cumulative number of entries behind the target wall over the first 5 entries during the probe trial (RM two-way ANOVA, p = 0.0484 effect of genotype). (**G**, **H**) KIBRA cKO does not affect overall movement as shown by equivalent total distance traveled (G) and moving velocity (H, velocity when mice are moving > 3cm/sec) during the probe trial (unpaired t-test, corrected for multiple comparisons). Total velocity including pauses was also not different between genotypes (WT, 18 ± 1cm/sec; cKO 16 ± 1cm/sec, ns). (**I**) KIBRA cKO and WT mice spend the same amount of time in the center of the maze during the probe trial (unpaired Mann-Whitney test). (**J**) No differences were observed between genotypes in weight lost due to time-restricted feeding (weight measured the last day of training/free feeding weight measured the day before maze habituation) (unpaired Mann-Whitney test). ***p < 0.001, **p < 0.01, *p < 0.05, 0.05 <#p < 0.1, n.s. p > 0.1.

### Acute reduction of KIBRA has no influence on long-term potentiation in the juvenile brain

Prior studies using a constitutive KIBRA KO model demonstrated no change in synaptic transmission or plasticity in juvenile mice (Makuch et al., 2011). It is possible that developmental compensation renders the juvenile brain resilient to KIBRA loss. In support of this notion, the KIBRA homolog WWC2, whose function in neurons is unknown, is selectively upregulated in juvenile constitutive KIBRA KO mice (Makuch et al., 2011). If constitutive deletion of KIBRA is compensated for by upregulation of other genes during early development, we hypothesized that rapid, acute deletion of KIBRA in juveniles may allow us to observe changes in synaptic function before genetic compensation occurs. Thus, to examine whether acute KIBRA deletion in juvenile mice impacts synaptic function, *Kibra^Fl/Fl^*:CaMKIICreER^T2^ mice were injected with tamoxifen between P14-P16 and synaptic transmission and plasticity were evaluated between P21-P25 (Fig. 6A). Using this system, KIBRA protein is reduced by approximately 60%, similar to the reduction observed in adult mice (% reduction in cKO mice, adult vs. juvenile, unpaired t-test, p = 0.332). We also demonstrate that Cre is not spontaneously active in juvenile CA1 (Fig. 6B-C). In contrast with the upregulation of WWC2 observed in constitutive KIBRA KO juvenile mice (Makuch et al., 2011), juvenile KIBRA cKO mice express normal levels of WWC2 (Fig. 6D) suggesting that long-term loss of KIBRA may produce compensatory upregulation of its homologs, but acute deletion does not.

We observe that acutely deleting KIBRA in CaMKIIα+ neurons of juvenile mice has no influence on the slope of the input-output curve across all experimental conditions (Fig. 7 A-C). In addition, we observe no effects on paired-pulse facilitation at CA1 synapses. Thus, as with adult neurons, acutely reducing KIBRA in the juvenile brain has no influence on the basal synaptic transmission. However, unlike adult neurons, acute reduction of KIBRA in juvenile mice had no impact on synaptic plasticity, supporting an age-specific role for KIBRA in synaptic function (Fig. 7E-F). Furthermore, these data suggest that compensatory upregulation of WWC2 as observed in constitutive KIBRA KO mice (Makuch et al., 2011) is unlikely to be the mechanism that confers resilience to loss of KIBRA in juvenile neurons.

**Figure 7.**
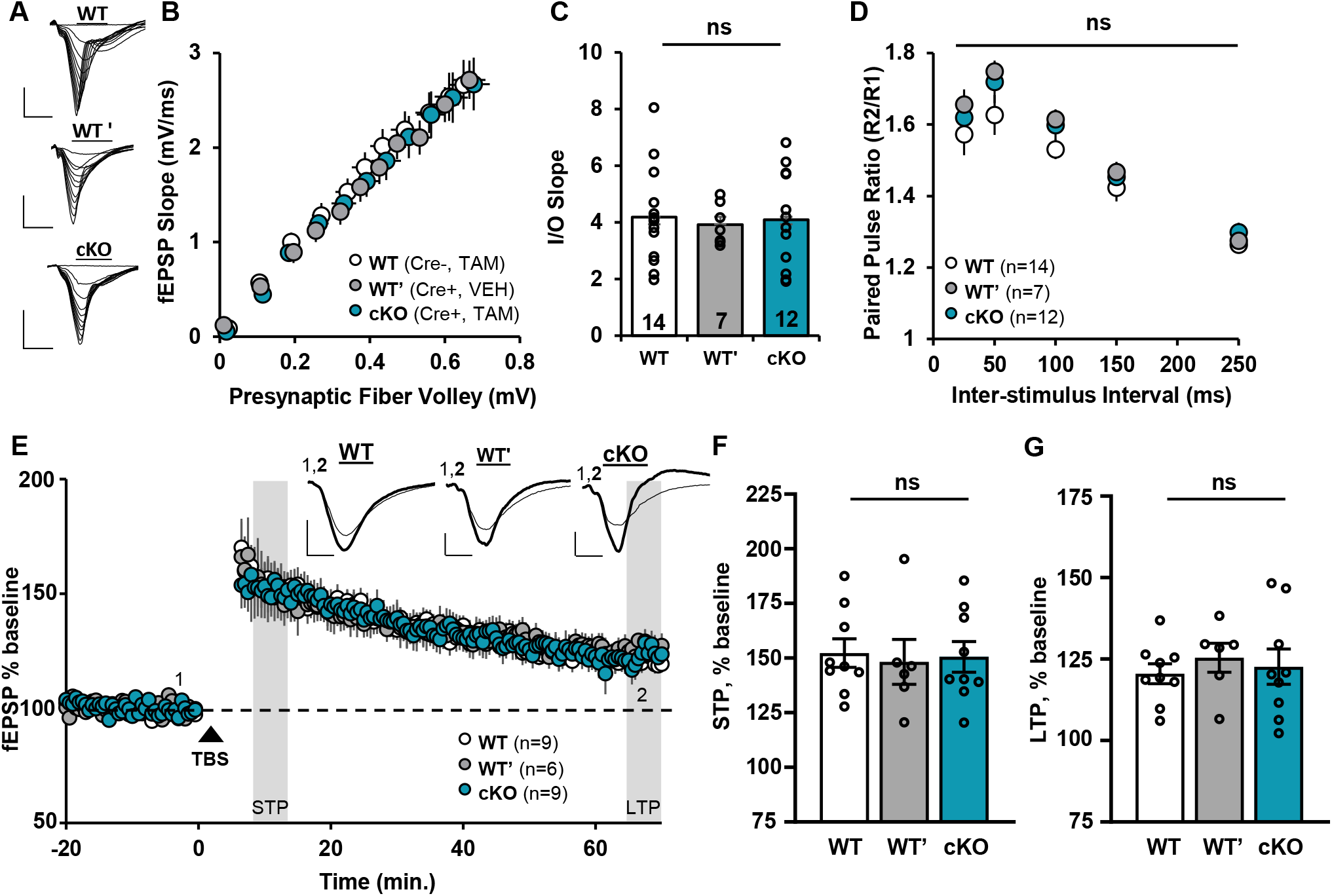
Acute reduction of KIBRA in the juvenile hippocampus does not affect basal synaptic transmission or long-term potentiation. (**A**) Representative traces from input-output curves, scale bars = 2mV/5ms, (**B**) Summary data from juvenile input-output analysis, (**C**) Slopes of individual input-output curves were quantified; no differences were observed across experimental conditions (one-way ANOVA, n.s.). Mean I-O slope: WT, 4.18 ± 0.45ms^-1^; WT’, 3.92 ± 0.27 ms-1; cKO, 4.09 ± 0.49 ms^-1^. (**D**) Paired pulse facilitation (fEPSP slope response 2/fEPSP slope response 1) is not altered in juvenile KIBRA cKO mice (RM two-way ANOVA, n.s. genotype X inter-stimulus interval interaction, main effect of genotype, and multiple comparisons at all inter-stimulus intervals). (**E**) Hippocampal LTP induced by four trains of theta-burst stimulation is unchanged after acute reduction of KIBRA in the juvenile brain. Scale bars= 0.25mV/5ms. (**F**) STP magnitude is unaffected in juvenile KIBRA cKO mice (one-way ANOVA, n.s.). 5min. avg at grey bar, WT, 152 ± 7 %; WT’, 148 ± 10 %; cKO, 151 ± 7%. (**G**) LTP magnitude measured at 65-70 mins post LTP induction is unaffected by acute reduction of KIBRA in the juvenile hippocampus (one-way ANOVA, n.s.). 5min. Avg at grey bar, WT, 120 ± 3 %; WT’, 125 ± 4 %; cKO, 123 ± 5%. All summary data presented as mean ± SEM. B-D: WT, n=14 slices from 4 mice; WT’, n = 7 slices from 2 mice; cKO, n = 12 slices from 4 mice. E-G: WT, n=9 slices from 4 mice; WT’, n = 8 slices from 2 mice; cKO, n = 9 slices from 4 mice.

### Acute reduction of KIBRA has minimal effects on the basal expression of AMPAR complexes in the juvenile brain

To determine if normal LTP in juvenile KIBRA cKO mice corresponded with intact pools of extrasynaptic AMPARs, we investigated how acute loss of KIBRA influences basal expression of AMPA receptors across subcellular fractions as described for adult mice. We observed no changes in total, membrane-associated, or synaptic GluA1 (Fig. 8).

**Figure 8.**
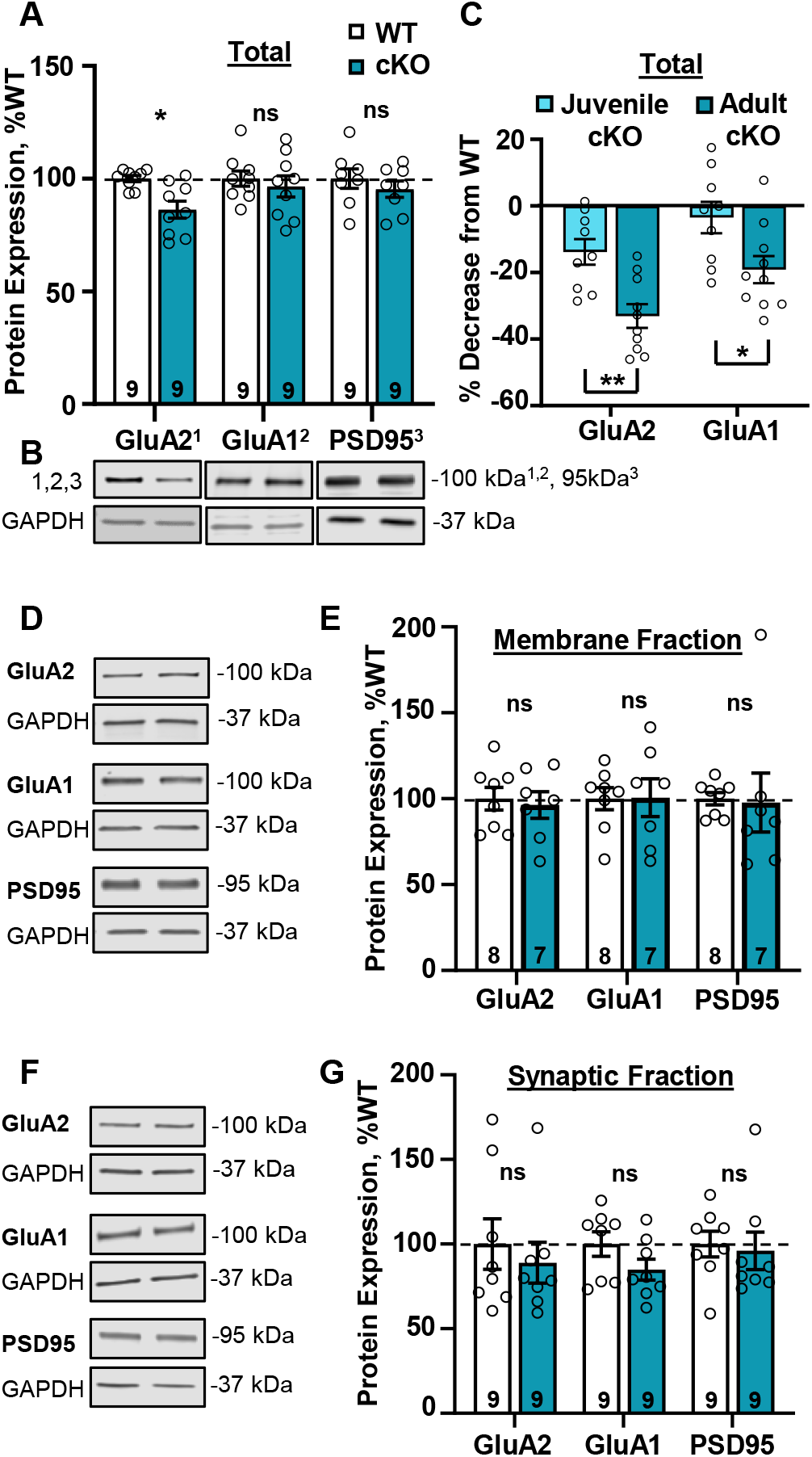
Acute reduction of KIBRA in the juvenile hippocampus has minimal effect on AMPAR expression. (**B,D,F)** Representative western blot images from sub-region CA1 of the juvenile hippocampus. (**A**) Acute reduction of KIBRA in the juvenile hippocampus decreases total expression of AMPAR subunit GluA2 but not GluA1 or PSD95 (unpaired t-tests, corrected for multiple comparisons, Welch’s correction for GluA2). GluA2, WT= 100 ± 1%, cKO=86 ±3; GluA1, WT=100 ± 3%, cKO=95 ± 4%; PSD95, WT=100 ± 4%, cKO=95 ± 3%. (**C**) Larger decrease in total AMPAR expression in adult compared to juvenile KIBRA cKO mice (unpaired t-tests, corrected for multiple comparisons). For each group, data is shown as % decrease from respective WT (GluA2, juvenile cKO = -14 ± 4%, adult cKO = -33 ± 4%; GluA1, juvenile cKO = -3 ± 5%, adult cKO = -19 ± 4%). (**E**) Acute reduction of KIBRA in the juvenile brain does not affect expression of membrane- associated AMPARs or PSD95 (unpaired t- (GluA1, GluA2) or Mann-Whitney (PSD95) tests, corrected for multiple comparisons). GluA2, WT=100 ± 6%, cKO=95 ± 4%; GluA1, WT=100 ± 6%, cKO=101 ± 11%; PSD95, WT=100 ± 3%, cKO=98 ± 17%. (**G**) Acute reduction of KIBRA in the juvenile brain does not alter basal expression of synaptic AMPA receptors or PSD95 (unpaired t- (GluA1, PSD95) or Mann-Whitney (GluA2) tests, corrected for multiple comparisons). GluA2, WT=100 ± 14%, cKO=89 ± 12%; GluA1, WT=100 ± 7%, cKO=85 ± 6%; PSD95, WT=100 ± 7%, cKO=96 ± 11%. Data shown as mean ± SEM, n on bar graphs = number of animals. *p < 0.05, **p < 0.01.

The excitatory synaptic scaffold PSD95 was similarly unaffected in juvenile KIBRA cKO mice. GluA2 expression was similarly unaffected in the membrane and synaptic fractions (Fig. 8D-G). Total GluA2 expression was modestly decreased in juvenile KIBRA cKO mice (Fig. 8A,B), but this decrease was significantly smaller than was observed in adult KIBRA cKO mice (Fig. 8C). Taken together, these data support the idea that dysregulation of extrasynaptic AMPAR pools contributes to impaired LTP in adult but not juvenile mice.

## Discussion

The post-synaptic scaffolding protein KIBRA is associated with normal variation in human memory performance (Papassotiropoulos et al., 2006; Almeida et al., 2008; Schaper et al., 2008; Bates et al., 2009; Preuschhof et al., 2009; Vassos et al., 2010; Yasuda et al., 2010; Kauppi et al., 2011; Pawlowski and Huentelman, 2011; Milnik et al., 2012; Duning et al., 2013; Muse et al., 2014; Vyas et al., 2014; Rovira et al., 2016).

Additionally, KIBRA and many of its binding partners are associated with childhood and adolescent-emergent NDDs (Hakak et al., 2001; Dev and Henley, 2006; Lauriat et al., 2006; Voineagu et al., 2011; Fromer et al., 2016; Hou et al., 2016; Kos et al., 2016; Parikshak et al., 2016; Willsey et al., 2017), highlighting KIBRA’s importance in cognitive function. However, the mechanisms through which KIBRA regulates adaptive cognition remain unclear. In addition, important questions remain regarding KIBRA’s acute role in adult neural function vs. its impact on neuronal and/or synaptic development. Prior work using constitutive deletion of KIBRA reported a role for KIBRA in plasticity in adult neurons (Makuch et al., 2011). However, these studies could not dissociate whether embryonic loss of KIBRA directly impacted synaptic plasticity in mature cells or instead produced earlier deficits in neuronal development which later manifested as impairments in plasticity. Here, using conditional, adult- or juvenile-specific deletion of KIBRA, we show that KIBRA plays an acute role in synaptic plasticity and maintenance of extrasynaptic pools of AMPARs selectively in adult neurons. We identify a novel role for KIBRA in LTP-induced upregulation of endogenous AMPAR expression and demonstrate that KIBRA function in CaMKIIα+ (predominantly excitatory) forebrain neurons is necessary for effective memory retention.

Synaptic AMPARs are responsible for the majority of fast excitatory neurotransmission in the central nervous system. Extrasynaptic AMPARs localized to the plasma membrane and in intracellular endosomes maintain essential reservoirs for activity-dependent increases in synaptic AMPAR content (Bourke et al., 2018; Parkinson and Hanley, 2018). Under basal conditions, AMPARs intrinsically recycle into and out of the synaptic plasma membrane. Internalized AMPARs destined for recycling back to the plasma membrane move from early to recycling endosomes, whereas receptors targeted for degradation are sorted into late endosomes for subsequent fusion with lysosomes (Widagdo et al., 2015; Bourke et al., 2018; Moretto and Passafaro, 2018; Parkinson and Hanley, 2018). Thus, a shift in the balance of endosomal trafficking of AMPARs toward lysosomes can alter AMPAR proteostasis and availability of these channels for synaptic insertion. We find that KIBRA is required to maintain extrasynaptic pools of AMPARs in adult neurons, consistent with prior studies demonstrating that KIBRA can stabilize its binding partners against degradation (Xiao et al., 2011; Vogt-Eisele et al., 2014; Hu et al., 2017; Ferguson et al., 2019; Song et al., 2019). Interestingly, we also see a reduction in the KIBRA homolog WWC2 in adult KIBRA cKO neurons. The function of WWC2 in neurons is unknown, so it is not clear if decreased WWC2 expression might contribute to LTP deficits in adult neurons.

One mechanism through which KIBRA may prevent AMPAR trafficking to lysosomes is through regulation of Rab family GTPases. Rab11 is enriched in recycling endosomes and disruption of Rab11 activity impairs AMPAR trafficking to the plasma membrane (Bowen et al., 2017; Bourke et al., 2018), consistent with our observation of decreased GluA2 in the membrane fraction following acute KIBRA deletion. KIBRA regulates trafficking through recycling endosomes in non-neuronal cells (Traer et al., 2007) and interacts with regulators of Rab11 activity in neurons (Fukuda et al., 2019). In non-neuronal cells, knockdown of KIBRA disrupts transport of transferrin receptors from early endosomes to Rab11-positive endocytic recycling compartment, resulting in lysosomal targeting and degradation of the receptors (Traer et al., 2007). In addition, KIBRA interacts with Rab27a (Song et al., 2019), and other components of the exocyst complex (Makuch et al., 2011) that facilitate trafficking of newly synthesized AMPARs to synapses (Gerges et al., 2006). Thus, our data support a model in which KIBRA directs endosomal AMPARs through recycling endosomes, away from late endosomes and lysosomal degradation. KIBRA also decreases ubiquitination of multiple binding partners (Xiao et al., 2011; Vogt-Eisele et al., 2014; Song et al., 2019) and AMPAR ubiquitination directs AMPAR trafficking toward lysosomes (Widagdo et al., 2015).

Thus, KIBRA may function through multiple pathways to maintain stable extrasynaptic pools of AMPARs and future studies should investigate the link between KIBRA and AMPAR ubiquitination.

Synaptic plasticity arises in distinct phases, each of which has unique mechanistic requirements. Upon LTP-inducing stimulation, extrasynaptic AMPARs residing in the plasma membrane are rapidly mobilized to synapses via lateral diffusion to support the early stages of LTP (STP) (Penn et al., 2017). We observe a strong trend toward decreased STP in adult KIBRA cKO mice with no change in charge transfer during TBS induction, consistent with depleted pools of extrasynaptic surface AMPARs under basal conditions in these mice. Maintenance of later phases of LTP requires *de novo* protein synthesis (Bradshaw et al., 2003; Abraham and Williams, 2008). AMPARs are synthesized in response to LTP stimuli (Nayak et al., 1998) and newly synthesized AMPARs are recruited to dendritic spines in neurons activated by learning (Matsuo et al., 2008). We find that a protein-synthesis-dependent form of LTP (Volk et al., 2013) induces rapid increases in AMPAR expression in WT, but not KIBRA cKO, neurons. Considering that we observe a decrease in basal AMPAR expression and given prior work demonstrating that KIBRA prevents degradation of its binding partners, we hypothesize that KIBRA acts to prevent rapid degradation of newly synthesized AMPARs, resulting in impaired late-phase LTP in adult KIBRA cKO mice. However, our data do not rule out a role for KIBRA in activity-induced protein synthesis (Deller et al., 2003; Duning et al., 2008).

We additionally confirmed that acute deletion of KIBRA in neurons of juvenile mice has no observable impact on synaptic plasticity, consistent with the lack of effect previously reported for juvenile mice with constitutive KIBRA deletion (Makuch et al., 2011).

Together, these findings strongly suggest that juvenile synapses can produce and maintain synaptic plasticity via a KIBRA-independent mechanism. These data do not necessarily rule out a role for KIBRA in juvenile plasticity, but rather highlight the fact that unlike adult neurons, immature neurons are able to compensate for KIBRA loss. Juvenile mice with constitutive KIBRA deletion display elevated expression of the KIBRA homolog WWC2 (Makuch et al., 2011), prompting the hypothesis that WWC2 may compensate for the loss of KIBRA in juvenile mice. However, in our conditional model in which KIBRA is acutely deleted, we do not observe a compensatory increase in WWC2. Thus, we conclude that WWC2 upregulation is not required for KIBRA- independent plasticity in juvenile neurons. Moreover, we demonstrate that, unlike in adult mice, extrasynaptic pools of AMPARs are largely intact in juvenile KIBRA cKO mice. Thus, normal plasticity in juvenile mice lacking KIBRA correlates with minimal changes in the maintenance of extrasynaptic AMPAR pools.

It is unclear why KIBRA deletion has no impact on immature synaptic plasticity. Interestingly, synapses from juvenile mice are also resilient to deletion of AMPAR subunits GluA1 or GluA2, as well as the AMPAR- and KIBRA-interacting protein PICK1, all of which produce impairments in plasticity when deleted from adult neurons (Jensen et al., 2003; Volk et al., 2010; Cao et al., 2018). Together, these findings point to a potential broader principle that plasticity in juvenile mice may utilize alternative or more robust mechanisms for LTP expression. In WT mice, KIBRA is strongly expressed at early developmental time points (Fig. 1A, (Johannsen et al., 2008)), arguing that KIBRA is likely playing an as-of-yet unidentified role in neuronal maturation. Indeed, neurons in juvenile KIBRA KO mice display morphological changes consistent with an early role for KIBRA that is distinct from its synaptic function in adult neurons (Blanque et al., 2015). While the observed decrease in total GluA2 expression is significantly larger in adult KIBRA cKO mice, we see a modest decrease in total GluA2 expression in juvenile mice, suggesting that AMPAR proteostasis may become KIBRA-dependent during adolescence.

KIBRA protein complexes are associated with several NDDs (Hakak et al., 2001; Dev and Henley, 2006; Lauriat et al., 2006; Voineagu et al., 2011; Fromer et al., 2016; Hou et al., 2016; Kos et al., 2016; Parikshak et al., 2016; Willsey et al., 2017). Recent genome-wide transcriptome analyses indicate that KIBRA is a developmentally regulated hub gene in ASD (Parikshak et al., 2016), highlighting the importance of identifying KIBRA’s function in the developing brain. Our work demonstrates that 21- to 25-day-old mice, an age roughly equivalent to a one-year-old human (Semple et al., 2013; Dutta and Sengupta, 2016), display normal synaptic function and plasticity in the absence of KIBRA. It will be important in future work to identify the time course in which KIBRA loss begins to impact excitatory synaptic plasticity, to examine non-synaptic functions of KIBRA during development, and to evaluate the role of KIBRA in other brain regions in which it is highly expressed (e.g. cortex).

In conclusion, we demonstrate an adult-selective role for KIBRA in maintaining proteostasis of extrasynaptic AMPARs under basal conditions, which correlates with impaired LTP in adult but not juvenile KIBRA cKO mice. We find that KIBRA in adult neurons is required for LTP-induced increases in endogenous AMPAR expression and for accurate maintenance of long-term memory, providing insight into the mechanisms by which this human memory-associated gene affects adaptive cognition.

## Materials and Methods

### Subjects

#### Mouse Breeding, Animal Husbandry, and Tamoxifen Treatment

All animals were housed in a climate-controlled environment on a 12 hours (hr) light/dark cycle. Food and water was provided ad libitum, with the exception of food restriction prior to Barnes Maze training (detailed below). All mice were bred on the background of Charles River C57Bl/6 mice (N10+). Inducible CaMKIIa-Cre^ERT2^ hemizygous mice were bred with homozygous Kibra *^floxed/floxed^(Makuch et al., 2011)* mice, yielding Cre positive (Cre +) and Cre negative (Cre -) Kibra *^floxed/floxed^* mice. To induce *Kibra* knockout, adult mice (2 to 4 months old) underwent five-days of tamoxifen or vehicle (sunflower seed oil) treatment consisting of 2 daily injections (IntraPeritoneal, 100mg/kg) separated by approximately 7 hours. Juvenile mice (P14) underwent three-days of tamoxifen treatment consisting of 1 daily injection of tamoxifen (IP 100 mg/kg) or vehicle. Both male and female mice were used throughout all experiments. All experiments were performed using protocols approved by the Institutional Animal Care and Use Committee at the University of Texas Southwestern Medical Center.

### Electrophysiology

#### Slice Preparation

Mice were anesthetized with Isoflurane prior to rapid decapitation. 380µm transverse hippocampal slices were prepared using a Leica VT 1200s vibratome following dissection of the hippocampus in ice cold oxygenated (95% O2/5% CO2) dissection buffer containing the following (in mM): 2.6 KCl, 1.25 NaH2PO4, 26 NaHCO3, 211 sucrose, 10 glucose, 0.75 CaCl2, 7MgCl2. Slices were recovered for at least 2hr in 30°C aCSF containing the following (in mM): 125 NaCl, 3.25 KCl, 25 NaHCO3,1.25 NaH2PO4·H2O, 11 glucose, 2CaCl2, 1MgCl2.

#### Ex Vivo Slice Electrophysiology

CA3 was removed prior to recording to prevent recurrent activity. Field excitatory postsynaptic potentials (fEPSPs) were digitally evoked (Cgynus Instruments, Model PG400A) at 0.033Hz with a 125µm platinum/ iridium concentric bipolar electrode (FHC, Bowdoinham, ME) placed in the stratum radiatum, approximately 50 μm below the stratum pyramidale. Glass recording electrodes filled with aCSF were positioned in the stratum radiatum ∼250μm away (orthodromic) from the stimulating electrode. Signals were amplified by a differential amplifier (Model 1800; A-M Systems), digitized using an Axon Instruments Digidata 1550A (Molecular Devices), and monitored using pClamp Clampex software (Molecular Devices). Recording aCSF and temperature were identical to recovery conditions, with a flow rate of ∼3mL/min. Input-output curves were obtained for each slice and responses were set to ∼45% max for LTP and paired-pulse ratio measurements. Paired-Pulse Ratios were recorded with inter-stimulus intervals of 25ms, 50ms, 100ms, 150ms, and 250ms.

*Theta Burst (TBS) LTP:* A stable baseline recording was obtained for a minimum of 20 minutes. LTP was then induced with TBS consisting of 4 trains separated by 10 seconds. Each train consisted of 10 bursts at 5Hz, with each burst containing 4 stimuli given at 100Hz.

#### TBS and Basal Stimulation for Quantifying LTP-induced AMPAR expression

Slice preparation and recording conditions were as described above, however two stimulating electrodes were used in order to cover most of stratum radiuatum (see Fig.4A): one stimulating electrode was placed ∼50μm below the stratum pyramidale in proximal CA1(orthodromic to recording electrode) and the other ∼120μm below the stratum pyramidale in distal CA1 (antidromic to recording electrode). Input-out curves were obtained for each slice and responses were set at ∼45% maximum. LTP: slices received brief (3-5min.) baseline stimulation (0.033Hz) prior to and following TBS. Baseline stimulation: slices received brief baseline stimulation (0.033Hz) for approximately 10 minutes. Following stimulation slices were gently removed from the recording chamber and returned to the recovery chamber (oxygenated aCSF at 30°C) for 30 or 120 minutes. CA1 was then micro-dissected under a Lecia S6e stereomicroscope and flash frozen in liquid nitrogen. Slices were processed for western blot by homogenization in boiling (100°C) SDS protein sample buffer containing; 15% glycerol, 94mM Tris/HCl pH 6.8, 3% sodium dodecyl sulfate (SDS), 0.02% bromophenol blue, 2% beta-mercaptoethanol. Each sample was homogenized for 15 seconds using an electric pestle and boiled for 10 minutes. 10µL of homogenate was analyzed for AMPAR expression via western blot as described below.

### Data Analysis

Electrophysiological data was analyzed using Clampfit 10.7. For LTP experiments, the fEPSP slope of individual responses was normalized to the average fEPSP slope from the 20 minute baseline immediately preceding TBS. LTP induction was assessed by measuring charge transfer (area under the curve) of each response from the end of the stimulation artifact + 9ms. For input-output curves, two responses were collected and averaged at each stimulation intensity. For PPR, four responses were collected and averaged at each inter-stimulus interval. Representative traces are averages of 4-8 individual responses, and stimulus artifacts have been removed for clarity.

### Molecular Analysis

### Hippocampal Dissection

Mice were briefly anaesthetized with isoflurane then rapidly decapitated. The brain was placed in ice cold dissection buffer (in mM: 125 NaCl, 3.25 KCl, 25 NaHCO3, 1.25 NaH2PO4·H2O, 11 glucose, 0.75 CaCl2, 7 MgCl2) and the hippocampus was removed, followed by isolation of area CA1 from the dentate gyrus and CA3. Immediately following dissection CA1 was flash frozen in liquid nitrogen and stored at -80 °C until processed as described below.

### Tissue Preparation for assessing hippocampal KIBRA expression

Isolated dorsal CA1 tissue (both hemispheres from one mouse) was homogenized in 300 microliters of homogenization buffer containing the following (in mM): 1% triton X- 100, 50 Tris-HCl, pH7.5, 150 NaCl, 0.2 okadaic acid, 1 NaPPi, 5 NaF, 1 NaVO3, and Roche Complete mini protease inhibitor cocktail. Tissue was homogenized by repetitively passing the tissue through a 26-gauge needle and syringe. Protein concentration was determined by using the Pierce^TM^ Detergent Compatible Bradford Assay Kit and a BioTek Synergy H1 Microplate Reader. Samples were boiled for 10 minutes in SDS protein sample buffer containing the following: 10% glycerol, 62.5mM Tris/HCl pH 6.8, 2% sodium dodecyl sulfate (SDS), 0.01% bromophenol blue, 1.25% beta-mercaptoethanol.

#### Subcellular Fractionation

Isolated CA1 tissue (both hemispheres from one mouse) was homogenized in 1 mL of homogenization buffer containing the following (in mM): 320 sucrose, 10 HEPES at pH 7.4, 1 EDTA, 0.2 okadaic acid, 1 NaPPi, 5 NaF, 1 NaVO3, Roche Complete Protease Inhibitor. 70 microliters of homogenate were removed for analysis of total protein expression. The remaining homogenate was then centrifuged at 800 xg for 10 minutes at 4°C. The supernatant (S1) was transferred to VWR high G-force micro-centrifuge tubes and centrifuged at 15,000 xg for 20 minutes at 4°C. The resultant supernatant (S2) was removed from the pellet (P2, membrane fraction), and the pellet was lysed in 500 microliters of Milli-Q water with protease and phosphatase inhibitors containing the following (in mM): 0.2 okadaic acid, 1 NaPPi, 5 NaF, 1 NaVO3, Roche Complete Protease Inhibitor. To ensure complete lysis of P2, 2 microliters of 1M HEPES at pH 7.4 was added to each sample followed by incubation with agitation for 30 minutes at 4°C. 70 microliters were removed from each sample following incubation and stored for future analysis of the membrane fraction. The remaining lysed P2 fraction was then centrifuged at 25,000 xg for 20 minutes at 4°C. The lysed supernatant (LS1) was removed, and the lysed pellet (LP1) was resuspended in 250 mL of 50mM HEPES pH 7.4 with protease and phosphatase inhibitors. Each sample was then rapidly mixed with an equal volume of 250μL of 1% triton X-100 with protease and phosphatase inhibitors and incubated with agitation at 4°C for 15 minutes. The resuspended LP1 was then centrifuged at 32,000 xg for 20 minutes at 4°C. The lysed supernatant (LS2) was discarded, thereby leaving the lysed pellet or PSD fraction. The PSD fraction was resuspended in 100μL of 50 mM HEPES pH 7.4 with proteasome and phosphatase inhibitors. Protein concentrations of each fraction were determined using the Pierce^TM^

Detergent Compatible Bradford Assay Kit. Samples were boiled for 10 minutes in SDS protein sample buffer containing the following: 10% glycerol, 62.5mM Tris/HCl pH 6.8, 2% sodium dodecyl sulfate (SDS), 0.01% bromophenol blue, 1.25% beta-mercaptoethanol.

#### Immunoblotting

To visualize the extent of the KIBRA KD, 45 μg of CA1 homogenate were loaded into 6% SDS-PAGE gels. Importantly, to verify that proteins were within a linear range for quantification, an in-gel protein concentration curve was performed by loading 50% (22.5 μg) and 150% (67.5 μg) of one sample. To ensure specificity of KIBRA Ab signal, tissue from a constitutive KIBRA knockout mouse was included, and showed no signal. Following separation, gels underwent a wet protein transfer to a PVDF membrane (Amersham Hybond P 0.45 PVDF) run in ice-cold transfer buffer at 100 V for 2 hours.

Membranes were blocked with a milk-based blocking solution containing the following: 1-2% milk and 0.5% TBST. The antibody conditions are outlined in table 2.1. The KIBRA signal was visualized using Amersham’s Highly Sensitive ECl Prime Reagent and the subsequent images were taken on a BioRad ChemiDoc. Images were taken every 30 seconds for a total exposure time of 30 minutes.

Hippocampal slices subjected to TBS/baseline stimulation and samples from the subcellular fractionation experiments used to examine AMPA receptor complex expression across the homogenate, plasma membrane, and synaptic fractions were loaded into either a 7% SDS-PAGE gel or 4-20% Gradient SDS- PAGE gel. Following separation, the gels underwent a wet protein transfer to a Nitrocellulose membrane (Odyssey Nitrocellulose Membrane, pore size 0.22 μm) run in ice- cold transfer buffer at 100 V for 2 hours. The membranes were then incubated in Odyssey Blocking Buffer (TBS) for 1 hour. Following blocking, the membranes were incubated in primary antibody overnight. Membranes were incubated with fluorescent secondary antibody for 1 hr and imaged on a Li-Cor Odyssey Scanner (Model 9120).

#### Antibodies

Conditions for each antibody were independently optimized to determine the appropriate amount of protein needed to achieve signal within linear range for quantification, determined by in-gel concentration curves as described above. 10 μg of protein was loaded for all homogenate western blots. 5 μg of protein was loaded for both the membrane and synaptic fractions, with the exception of blots used for examining PSD95, in which 2 μg of protein was loaded for the membrane and synaptic fractions.

Refer to table 2.1 for the full list of antibody concentrations used.

#### Data Analysis

Western blot data was quantified using the Li-Cor image studio lite software. Each sample was normalized to a loading control before being normalized to the average of all WT (Cre-negative tamoxifen-treated) samples within the same gel.

### Behavior

#### Apparatus

The modified Barnes Maze consisted of a 96cm diameter white circular platform with 11 equally spaced walls. Each wall was 13cm tall x 14cm wide, placed 8.5cm from the outside edge of the maze, and had a triangular back that allowed mice to explore behind the wall but prevented mice from traversing the maze by walking around the outside edge behind the walls (see Fig. 5A). During training sessions, a covered target box (10.5cm long, 9cm wide, 9.5cm tall) was located behind one of the walls, and was not visible until the mouse walked behind the wall. ∼0.5cm^3^ pieces of peanut butter chip were placed at the back of the target box during training. Additionally, a physically inaccessible peanut butter chip was placed in the center of each wall barrier to prevent mice from using the smell of the peanut butter chip to navigate to the target box. The target box location relative to extra-maze room cues remained the same for all training sessions and probe trials. The start chamber was a 9cm cup with opaque sides. The maze was located 42cm above the floor in a brightly lit dedicated behavior room.

#### Food restriction and animal handling

Mice were handled 5 minutes per day for 5 days immediately prior to training. During the handling period, mice were introduced to peanut butter chips (one Reeses peanut butter chip per mouse each day). During training and on the day prior to the probe trial, mice underwent time-restricted feeding during which they were allowed ad lib access to food for two hours (∼6pm -8pm) per day.

#### Training and Testing

Mice remained group housed throughout behavioral training and testing. Behavior was conducted during the light cycle, between 9am and 5pm.

Day 0, habituation: Mice were placed in the center of the arena in an opaque start chamber. Mice were released from the start chamber 5 seconds after entering the arena, and were gently guided to the target box, and the entrance to the box was blocked. Mice remained in the target box for 2 minutes before being returned to their home cage.

Days 1-4, training: Mice were placed in the center of the arena in an opaque start chamber. Mice were released from the start chamber 5 seconds after entering the arena. Because mice can turn freely in the start chamber, mice began each trial facing a random direction. Mice were allowed 90 seconds to find the target box. Mice that did not enter within this time were gently guided to the target box by the experimenter.

Mice remained in the closed target box for 30 seconds before being returned to the home cage. Mice underwent four training trials per day, with an inter-trial interval of 10 minutes. Mice were returned to holding cages between trials. The maze was cleaned with water and rotated after each trial to prevent mice from using scent cues to navigate to the target box. The target box was moved after each maze rotation such that it remained in the same location relative to the room cues.

Probe trial: The probe trial, conducted 7 days after the last training session, was identical to a training trial except that the target box was removed.

#### Data Analysis

All training sessions and probe trials were continuously recorded on a SONY HDR- CX440 digital video camera. Videos were sampled at 30 FPS with a resolution of at least 1280x720. Mouse position was obtained using DeepLabCut (Mathis et al., 2018) to train a deep convolutional network. Briefly, a set of training frames were extracted from a representative subset of videos throughout the behavioral paradigm using a k means algorithm. Training frames were then manually labeled with the location of the animal’s body parts (head, snout, ears, and tail). After training, the model was evaluated using a train-test split to compute prediction error. Optimization was done through active refinement of poorly predicted frames and retraining to maintain high prediction accuracy across days and sessions. A final review of position estimation results was done using labeled videos to assess generalization across animals. The position data generated by Deeplabcut was then analyzed with custom MATLAB (MathWorks) programs, code available on request. The number of entries behind each wall was scored by an experimenter blind to genotype (‘entry’ entailed the mouse’s entire body crossing the plane of the walls) and confirmed by automated MATLAB analysis of DeepLabCut-generated positional data. To calculate zone occupancy during the probe trial, the maze was divided into a 56cm diameter central zone, and the outer portion of the maze was divided into 11 equally sized sections centered each gap leading behind a wall. The target zone consisted of the section previously attached to the target box during training plus the sections on either side, the ‘right’ zone contained the three sections clockwise from the target zone, ‘opposite’ contained the three sections opposite the target zone, and ‘left’ contained the two sections counterclockwise from the target zone. Zone occupancy was normalized for the number of sections in the zone.

#### Statistics

Statistical analysis was performed in Graph Pad Prism version 9.3.0. Data are plotted as mean ± SEM unless otherwise noted. The statistical test performed for each analysis is noted in the figure legend. p < 0.05 was considered significant. Data and test residuals were assessed for normality using Shapiro-wilk and D’Agostino-Pearson tests, in addition to visual inspection of QQ plots. Equality of variance between groups was assessed using an F test (t-tests) or Brown-Forsythe test (ANOVA), and examination of residual homoscedasticity plots. For data that did not meet criteria for equal variance a Welch’s correction was applied to the statistical test. Sphericity (equal variability of differences) was not assumed for repeated measures ANOVAs (Geisser-Greenhouse correction was applied). One or two- way ANOVAs with post hoc multiple comparison tests as needed or t-tests were used to evaluate significance. The Holm-Šidák correction was applied to all multiple comparisons (ANOVAs and multiple t-tests). Statistical tests were chosen based on sample size, hypotheses, and agreement with statistical assumptions. In the small number of instances where assumptions were not met for the above tests, nonparametric statistics were used (noted in figure legends).

## Acknowledgements

This work was supported by National Institutes of Health Grant NIMH 1R01MH117149-01. M.L.M. was supported by the Howard Hughes Medical Institute Gilliam Fellowship for Advanced Study. L.Q. is supported by National Institutes of Health Grant 1F99NS120543-01. We thank Dr. Brad Pfeiffer for manuscript feedback and for providing Matlab programs.

## Notes

**Conflict of Interest:** The authors declare no competing financial interests.

### Competing Interest Statement

The authors have declared no competing interest.

## References

Abraham WC, Williams JM (2008) LTP maintenance and its protein synthesis-dependence. Neurobiology of learning and memory 89:260–268.

Alberini CM, Travaglia A (2017) Infantile Amnesia: A Critical Period of Learning to Learn and Remember. The Journal of neuroscience : the official journal of the Society for Neuroscience 37:5783–5795.

Allen-Institute_Brain-Atlas © 2004 Allen Institute for Brain Science. Allen Mouse Brain Atlas. Available from: https://mouse.brain-map.org/.

Allen-Institute_Cell-Types © 2015 Allen Institute for Brain Science. Allen Cell Types Database (2015). Available from: https://celltypes.brain-map.org/rnaseq/mouse/v1-alm.

Almeida OP, Schwab SG, Lautenschlager NT, Morar B, Greenop KR, Flicker L, Wildenauer D (2008) KIBRA genetic polymorphism influences episodic memory in later life, but does not increase the risk of mild cognitive impairment. J Cell Mol Med 12:1672–1676.

Bates TC, Price JF, Harris SE, Marioni RE, Fowkes FG, Stewart MC, Murray GD, Whalley LJ, Starr JM, Deary IJ (2009) Association of KIBRA and memory. Neurosci Lett 458:140–143.

Blanque A, Repetto D, Rohlmann A, Brockhaus J, Duning K, Pavenstadt H, Wolff I, Missler M (2015) Deletion of KIBRA, protein expressed in kidney and brain, increases filopodial-like long dendritic spines in neocortical and hippocampal neurons in vivo and in vitro. Frontiers in neuroanatomy 9:13.

Bourke AM, Bowen AB, Kennedy MJ (2018) New approaches for solving old problems in neuronal protein trafficking. Molecular and cellular neurosciences 91:48–66.

Bowen AB, Bourke AM, Hiester BG, Hanus C, Kennedy MJ (2017) Golgi-independent secretory trafficking through recycling endosomes in neuronal dendrites and spines. eLife 6.

Bradshaw KD, Emptage NJ, Bliss TV (2003) A role for dendritic protein synthesis in hippocampal late LTP. The European journal of neuroscience 18:3150–3152.

Buther K, Plaas C, Barnekow A, Kremerskothen J (2004) KIBRA is a novel substrate for protein kinase Czeta. Biochem Biophys Res Commun 317:703–707.

Cao F, Zhou Z, Cai S, Xie W, Jia Z (2018) Hippocampal Long-Term Depression in the Presence of Calcium-Permeable AMPA Receptors. Frontiers in synaptic neuroscience 10:41.

Cembrowski MS, Wang L, Sugino K, Shields BC, Spruston N (2016) Hipposeq: a comprehensive RNA-seq database of gene expression in hippocampal principal neurons. eLife 5:e14997.

Chiu SL, Diering GH, Ye B, Takamiya K, Chen CM, Jiang Y, Niranjan T, Schwartz CE, Wang T, Huganir RL (2017) GRASP1 Regulates Synaptic Plasticity and Learning through Endosomal Recycling of AMPA Receptors. Neuron 93:1405–1419.e1408.

Choquet D (2018) Linking Nanoscale Dynamics of AMPA Receptor Organization to Plasticity of Excitatory Synapses and Learning. The Journal of neuroscience : the official journal of the Society for Neuroscience 38:9318–9329.

Deller T, Korte M, Chabanis S, Drakew A, Schwegler H, Stefani GG, Zuniga A, Schwarz K, Bonhoeffer T, Zeller R, Frotscher M, Mundel P (2003) Synaptopodin-deficient mice lack a spine apparatus and show deficits in synaptic plasticity. Proceedings of the National Academy of Sciences of the United States of America 100:10494–10499.

Dev KK, Henley JM (2006) The schizophrenic faces of PICK1. Trends in Pharmacological Sciences 27:574–579.

Diering GH, Huganir RL (2018) The AMPA Receptor Code of Synaptic Plasticity. Neuron 100:314–329.

Dumas TC (2005) Late postnatal maturation of excitatory synaptic transmission permits adult-like expression of hippocampal-dependent behaviors. Hippocampus 15:562–578.

Duning K, Schurek EM, Schluter M, Bayer M, Reinhardt HC, Schwab A, Schaefer L, Benzing T, Schermer B, Saleem MA, Huber TB, Bachmann S, Kremerskothen J, Weide T, Pavenstadt H (2008) KIBRA modulates directional migration of podocytes. Journal of the American Society of Nephrology : JASN 19:1891–1903.

Duning K et al. (2013) Common exonic missense variants in the C2 domain of the human KIBRA protein modify lipid binding and cognitive performance. Translational psychiatry 3:e272.

Dutta S, Sengupta P (2016) Men and mice: Relating their ages. Life Sci 152:244–248.

Erdmann G, Schutz G, Berger S (2007) Inducible gene inactivation in neurons of the adult mouse forebrain. BMC Neuroscience 8:63.

Farooq U, Dragoi G (2019) Emergence of preconfigured and plastic time-compressed sequences in early postnatal development. Science 363:168–173.

Ferguson L, Hu J, Cai D, Chen S, Dunn TW, Pearce K, Glanzman DL, Schacher S, Sossin WS (2019) Isoform Specificity of PKMs during Long-Term Facilitation in Aplysia Is Mediated through Stabilization by KIBRA. The Journal of neuroscience : the official journal of the Society for Neuroscience 39:8632–8644.

Fromer M et al. (2016) Gene expression elucidates functional impact of polygenic risk for schizophrenia. Nat Neurosci 19:1442–1453.

Fukuda T, Nagashima S, Inatome R, Yanagi S (2019) CAMDI interacts with the human memory-associated protein KIBRA and regulates AMPAR cell surface expression and cognition. PLoS One 14:e0224967.

Gerges NZ, Backos DS, Rupasinghe CN, Spaller MR, Esteban JA (2006) Dual role of the exocyst in AMPA receptor targeting and insertion into the postsynaptic membrane. The EMBO journal 25:1623–1634.

Habib N, Li Y, Heidenreich M, Swiech L, Avraham-Davidi I, Trombetta JJ, Hession C, Zhang F, Regev A (2016) Div-Seq: Single-nucleus RNA-Seq reveals dynamics of rare adult newborn neurons. Science 353:925–928.

Hakak Y, Walker JR, Li C, Wong WH, Davis KL, Buxbaum JD, Haroutunian V, Fienberg AA (2001) Genome-wide expression analysis reveals dysregulation of myelination-related genes in chronic schizophrenia. Proceedings of the National Academy of Sciences of the United States of America 98:4746–4751.

Heitz FD, Farinelli M, Mohanna S, Kahn M, Duning K, Frey MC, Pavenstadt H, Mansuy IM (2016) The memory gene KIBRA is a bidirectional regulator of synaptic and structural plasticity in the adult brain. Neurobiology of learning and memory 135:100–114.

Hou L et al. (2016) Genome-wide association study of 40,000 individuals identifies two novel loci associated with bipolar disorder. Human molecular genetics 25:3383–3394.

Hu J, Ferguson L, Adler K, Farah CA, Hastings MH, Sossin WS, Schacher S (2017) Selective Erasure of Distinct Forms of Long-Term Synaptic Plasticity Underlying Different Forms of Memory in the Same Postsynaptic Neuron. Current biology : CB 27:1888–1899.e1884.

Huganir RL, Nicoll RA (2013) AMPARs and synaptic plasticity: the last 25 years. Neuron 80:704–717.

Jensen V, Kaiser KM, Borchardt T, Adelmann G, Rozov A, Burnashev N, Brix C, Frotscher M, Andersen P, Hvalby Ø, Sakmann B, Seeburg PH, Sprengel R (2003) A juvenile form of postsynaptic hippocampal long-term potentiation in mice deficient for the AMPA receptor subunit GluR-A. The Journal of physiology 553:843–856.

Johannsen S, Duning K, Pavenstadt H, Kremerskothen J, Boeckers TM (2008) Temporal-spatial expression and novel biochemical properties of the memory-related protein KIBRA. Neuroscience 155:1165–1173.

Kauppi K, Nilsson LG, Adolfsson R, Eriksson E, Nyberg L (2011) KIBRA polymorphism is related to enhanced memory and elevated hippocampal processing. The Journal of neuroscience : the official journal of the Society for Neuroscience 31:14218–14222.

Kos MZ, Carless MA, Peralta J, Blackburn A, Almeida M, Roalf D, Pogue-Geile MF, Prasad K, Gur RC, Nimgaonkar V, Curran JE, Duggirala R, Glahn DC, Blangero J, Gur RE, Almasy L (2016) Exome Sequence Data From Multigenerational Families Implicate AMPA Receptor Trafficking in Neurocognitive Impairment and Schizophrenia Risk. Schizophr Bull 42:288–300.

Kremerskothen J, Kindler S, Finger I, Veltel S, Barnekow A (2006) Postsynaptic recruitment of Dendrin depends on both dendritic mRNA transport and synaptic anchoring. Journal of neurochemistry 96:1659–1666.

Kremerskothen J, Plaas C, Buther K, Finger I, Veltel S, Matanis T, Liedtke T, Barnekow A (2003) Characterization of KIBRA, a novel WW domain-containing protein. Biochem Biophys Res Commun 300:862–867.

Lauriat TL, Dracheva S, Kremerskothen J, Duning K, Haroutunian V, Buxbaum JD, Hyde TM, Kleinman JE, McInnes LA (2006) Characterization of KIAA0513, a novel signaling molecule that interacts with modulators of neuroplasticity, apoptosis, and the cytoskeleton. Brain Res 1121:1–11.

Lohmann C, Kessels HW (2014) The developmental stages of synaptic plasticity. The Journal of physiology 592:13–31.

Makuch L, Volk L, Anggono V, Johnson RC, Yu Y, Duning K, Kremerskothen J, Xia J, Takamiya K, Huganir RL (2011) Regulation of AMPA receptor function by the human memory-associated gene KIBRA. Neuron 71:1022–1029.

Mathis A, Mamidanna P, Cury KM, Abe T, Murthy VN, Mathis MW, Bethge M (2018) DeepLabCut: markerless pose estimation of user-defined body parts with deep learning. Nat Neurosci 21:1281–1289.

Matsuo N, Reijmers L, Mayford M (2008) Spine-type-specific recruitment of newly synthesized AMPA receptors with learning. Science 319:1104–1107.

Milnik A, Heck A, Vogler C, Heinze HJ, de Quervain DJ, Papassotiropoulos A (2012) Association of KIBRA with episodic and working memory: a meta-analysis. American journal of medical genetics Part B, Neuropsychiatric genetics : the official publication of the International Society of Psychiatric Genetics 159b:958–969.

Moretto E, Passafaro M (2018) Recent Findings on AMPA Receptor Recycling. Frontiers in cellular neuroscience 12:286.

Muse J, Emery M, Sambataro F, Lemaitre H, Tan HY, Chen Q, Kolachana BS, Das S, Callicott JH, Weinberger DR, Mattay VS (2014) WWC1 genotype modulates age-related decline in episodic memory function across the adult life span. Biological psychiatry 75:693–700.

Nayak A, Zastrow DJ, Lickteig R, Zahniser NR, Browning MD (1998) Maintenance of late-phase LTP is accompanied by PKA-dependent increase in AMPA receptor synthesis. Nature 394:680–683.

Papassotiropoulos A, Stephan DA, Huentelman MJ, Hoerndli FJ, Craig DW, Pearson JV, Huynh KD, Brunner F, Corneveaux J, Osborne D, Wollmer MA, Aerni A, Coluccia D, Hanggi J, Mondadori CR, Buchmann A, Reiman EM, Caselli RJ, Henke K, de Quervain DJ (2006) Common Kibra alleles are associated with human memory performance. Science 314:475–478.

Parikshak NN, Swarup V, Belgard TG, Irimia M, Ramaswami G, Gandal MJ, Hartl C, Leppa V, Ubieta LT, Huang J, Lowe JK, Blencowe BJ, Horvath S, Geschwind DH (2016) Genome-wide changes in lncRNA, splicing, and regional gene expression patterns in autism. Nature 540:423–427.

Parkinson GT, Hanley JG (2018) Mechanisms of AMPA Receptor Endosomal Sorting. Frontiers in molecular neuroscience 11:440.

Pawlowski TL, Huentelman MJ (2011) Identification of a common variant affecting human episodic memory performance using a pooled genome-wide association approach: a case study of disease gene identification. Methods in molecular biology (Clifton, NJ) 700:261–269.

Penn AC, Zhang CL, Georges F, Royer L, Breillat C, Hosy E, Petersen JD, Humeau Y, Choquet D (2017) Hippocampal LTP and contextual learning require surface diffusion of AMPA receptors. Nature 549:384–388.

Preuschhof C, Heekeren HR, Li SC, Sander T, Lindenberger U, Backman L (2009) KIBRA and CLSTN2 polymorphisms exert interactive effects on human episodic memory. Neuropsychologia.

Rocca DL, Martin S, Jenkins EL, Hanley JG (2008) Inhibition of Arp2/3-mediated actin polymerization by PICK1 regulates neuronal morphology and AMPA receptor endocytosis. Nature cell biology 10:259–271.

Rosse C, Formstecher E, Boeckeler K, Zhao Y, Kremerskothen J, White MD, Camonis JH, Parker PJ (2009) An aPKC-exocyst complex controls paxillin phosphorylation and migration through localised JNK1 activation. PLoS Biol 7:e1000235.

Rovira E, Mackie RS, Clark N, Squire PN, Hendricks MD, Pulido AM, Greenwood PM (2016) A role for attention during wilderness navigation: Comparing effects of BDNF, KIBRA, and CHRNA4. Neuropsychology 30:709–719.

Schaper K, Kolsch H, Popp J, Wagner M, Jessen F (2008) KIBRA gene variants are associated with episodic memory in healthy elderly. Neurobiology of aging 29:1123–1125.

Semple BD, Blomgren K, Gimlin K, Ferriero DM, Noble-Haeusslein LJ (2013) Brain development in rodents and humans: Identifying benchmarks of maturation and vulnerability to injury across species. Progress in neurobiology 106–107:1-16.

Shepherd JD, Huganir RL (2007) The Cell Biology of Synaptic Plasticity: AMPA Receptor Trafficking. Annu Rev Cell Dev Biol 23:613–643.

Song L, Tang S, Han X, Jiang Z, Dong L, Liu C, Liang X, Dong J, Qiu C, Wang Y, Du Y (2019) KIBRA controls exosome secretion via inhibiting the proteasomal degradation of Rab27a. Nature communications 10:1639.

Tracy Tara E, Sohn Peter D, Minami SS, Wang C, Min S-W, Li Y, Zhou Y, Le D, Lo I, Ponnusamy R, Cong X, Schilling B, Ellerby Lisa M, Huganir Richard L, Gan L (2016) Acetylated Tau Obstructs KIBRA-Mediated Signaling in Synaptic Plasticity and Promotes Tauopathy-Related Memory Loss. Neuron 90:245–260.

Traer CJ, Rutherford AC, Palmer KJ, Wassmer T, Oakley J, Attar N, Carlton JG, Kremerskothen J, Stephens DJ, Cullen PJ (2007) SNX4 coordinates endosomal sorting of TfnR with dynein-mediated transport into the endocytic recycling compartment. Nature cell biology 9:1370–1380.

Tushev G, Glock C, Heumüller M, Biever A, Jovanovic M, Schuman EM (2018) Alternative 3’ UTRs Modify the Localization, Regulatory Potential, Stability, and Plasticity of mRNAs in Neuronal Compartments. Neuron 98:495–511.e496.

Vassos E, Bramon E, Picchioni M, Walshe M, Filbey FM, Kravariti E, McDonald C, Murray RM, Collier DA, Toulopoulou T (2010) Evidence of association of KIBRA genotype with episodic memory in families of psychotic patients and controls. Journal of psychiatric research 44:795–798.

Vogt-Eisele A, Kruger C, Duning K, Weber D, Spoelgen R, Pitzer C, Plaas C, Eisenhardt G, Meyer A, Vogt G, Krieger M, Handwerker E, Wennmann DO, Weide T, Skryabin BV, Klugmann M, Pavenstadt H, Huentelmann MJ, Kremerskothen J, Schneider A (2014) KIBRA (KIdney/BRAin protein) regulates learning and memory and stabilizes Protein kinase Mzeta. Journal of neurochemistry 128:686–700.

Voineagu I, Wang X, Johnston P, Lowe JK, Tian Y, Horvath S, Mill J, Cantor RM, Blencowe BJ, Geschwind DH (2011) Transcriptomic analysis of autistic brain reveals convergent molecular pathology. Nature 474:380–384.

Volk L, Kim C-H, Takamiya K, Yu Y, Huganir RL (2010) Developmental regulation of protein interacting with C kinase 1 (PICK1) function in hippocampal synaptic plasticity and learning. Proceedings of the National Academy of Sciences 107:21784–21789.

Volk LJ, Bachman JL, Johnson R, Yu Y, Huganir RL (2013) PKM-zeta is not required for hippocampal synaptic plasticity, learning and memory. Nature 493:420–423.

Vyas NS, Ahn K, Stahl DR, Caviston P, Simic M, Netherwood S, Puri BK, Lee Y, Aitchison KJ (2014) Association of KIBRA rs17070145 polymorphism with episodic memory in the early stages of a human neurodevelopmental disorder. Psychiatry research 220:37–43.

Widagdo J, Chai YJ, Ridder MC, Chau YQ, Johnson RC, Sah P, Huganir RL, Anggono V (2015) Activity-Dependent Ubiquitination of GluA1 and GluA2 Regulates AMPA Receptor Intracellular Sorting and Degradation. Cell reports 10:783–795.

Willsey AJ, Fernandez TV, Yu D, King RA, Dietrich A, Xing J, Sanders SJ, Mandell JD, Huang AY, Richer P, Smith L, Dong S, Samocha KE, Neale BM, Coppola G, Mathews CA, Tischfield JA, Scharf JM, State MW, Heiman GA (2017) De Novo Coding Variants Are Strongly Associated with Tourette Disorder. Neuron 94:486–499.e489.

Xiao L, Chen Y, Ji M, Dong J (2011) KIBRA regulates Hippo signaling activity via interactions with large tumor suppressor kinases. The Journal of biological chemistry 286:7788–7796.

Yasuda Y, Hashimoto R, Ohi K, Fukumoto M, Takamura H, Iike N, Yoshida T, Hayashi N, Takahashi H, Yamamori H, Morihara T, Tagami S, Okochi M, Tanaka T, Kudo T, Kamino K, Ishii R, Iwase M, Kazui H, Takeda M (2010) Association study of KIBRA gene with memory performance in a Japanese population. The world journal of biological psychiatry : the official journal of the World Federation of Societies of Biological Psychiatry 11:852–857.

Zhang L, Yang S, Wennmann DO, Chen Y, Kremerskothen J, Dong J (2014) KIBRA: In the brain and beyond. Cellular signalling 26:1392–1399.

